# Gut barrier defects, increased intestinal innate immune response, and enhanced lipid catabolism drive lethality in *N*-glycanase 1 deficient *Drosophila*

**DOI:** 10.1101/2023.04.07.536022

**Authors:** Ashutosh Pandey, Antonio Galeone, Seung Yeop Han, Benjamin A Story, Gaia Consonni, William F Mueller, Lars M Steinmetz, Thomas Vaccari, Hamed Jafar-Nejad

## Abstract

Intestinal barrier dysfunction leads to inflammation and associated metabolic changes. However, the relative impact of infectious versus non-infectious mechanisms on animal health in the context of barrier dysfunction is not well understood. Here, we establish that loss of *Drosophila N*-glycanase 1 (Pngl) leads to gut barrier defects, which cause starvation and increased JNK activity. These defects result in Foxo overactivation, which induces a hyperactive innate immune response and lipid catabolism, thereby contributing to lethality associated with loss of *Pngl*. Notably, germ-free rearing of *Pngl* mutants did not rescue lethality. In contrast, raising *Pngl* mutants on isocaloric, fat-rich diets improved animal survival in a dosage-dependent manner. Our data indicate that Pngl functions in *Drosophila* larvae to establish the gut barrier, and that the immune and metabolic consequences of loss of *Pngl* are primarily mediated through non-infectious mechanisms.

## Introduction

Intestinal barrier dysfunction allows various pathogens and non-infectious stimuli to induce the innate immune response^1–5^. Although the goal of innate immune response induction is to restore intestinal homeostasis, its hyperactivation can have local and systemic adverse effects and is implicated in the pathogenesis of human diseases including autoimmune and neurodegenerative disorders^6–8^. In addition, hyperactive immune response can be accompanied by profound changes in metabolism including high energy demand and subsequent depletion of the nutrient depot^9,10^. For example, induction of chronic inflammation results in the aggravation of metabolic disorders in mice models of obesity^11^. However, the relative contribution of infectious versus non-infectious mechanisms to detrimental consequences caused by gut barrier dysfunction is not well understood.

The gut mucus layer is one of the key components regulating intestinal barrier functions^12^. Equivalent to the mammalian gut mucus layer, a peritrophic matrix (PM) is present in the *Drosophila* intestine^13^. The PM is composed of highly glycosylated proteins and chitin, and is continuously secreted from a group of specialized cells called <u>p</u>eritrophic matrix-forming ring (PR) cells in the proventriculus region at the junction of foregut and midgut^13,14^. There is strong evidence that in addition to chitins and mucin-type *O-*glycans, PM also contains *N*-glycoproteins^15,16^. Moreover, lectin-based studies in another insect suggest that *N*-glycoproteins might control the functional properties of PM like its permeability^17^. However, genetic evidence for the contribution of *N*-glycoproteins to gut barrier function in *Drosophila* is lacking.

The deglycosylating enzyme *N*-glycanase 1 (NGLY1) removes *N*-glycan from misfolded proteins and is thought to function in the endoplasmic reticulum-associated degradation (ERAD) pathway^18–20^. Recessive mutations in human *NGLY1* cause a congenital disorder of deglycosylation named NGLY1 deficiency^21–23^. It is an ultra-rare disorder that leads to global developmental delay and affects multiple organ systems including the nervous system and the gastrointestinal system. Loss of the *Drosophila* homolog of human NGLY1 (PNGase-like or Pngl) results in semi-lethality, as less than 1% homozygous mutant animals finish the larval and pupal development and eclose as an adult organism^24,25^. We have previously reported that loss of *Pngl* in the visceral mesoderm impairs signaling pathways mediated by <u>d</u>eca<u>p</u>enta<u>p</u>legic (Dpp; homolog of human bone morphogenetic protein 4, BMP4) and <u>a</u>denosine <u>m</u>ono<u>p</u>hosphate-activated protein <u>k</u>inase (AMPK) in the larval intestine, which lead to structural and functional intestinal phenotypes and contribute to the lethality of *Pngl^–/–^* animals^25,26^. However, impaired BMP and AMPK signaling due to mesodermal loss of *Pngl* did not fully explain the lethality of *Pngl* mutants, suggesting critical roles for Pngl in other biological processes and potentially in other cell types.

Here, we report that loss of *Pngl* leads to increased expression of the innate immune genes in the *Drosophila* larval intestine and a systemic increase in lipid catabolism, compromising developmental progression and leading to lethality. We find that loss of *Pngl* causes intracellular accumulation of *N*-glycoproteins in the PR cells, accompanied by abnormalities in peritrophic matrix and impairment in the gut barrier function. Our data suggest that the gut barrier defects result in increased activation of Foxo in the gut epithelial cells, both via enhanced stress-induced JNK signaling and through a systemic starvation response. In addition, we observe that Pngl is required cell-autonomously in enterocytes and in the fat body to prevent aberrant Foxo activation and to repress lipid catabolism. Importantly, while germ-free rearing does not rescue the lethality of *Pngl* mutants, increasing the lipid content in isocaloric diets improves their survival to adulthood, suggesting that the mutant animals lack sufficient energy stores to reach the adult stage. Altogether, our data suggest that Pngl is required to establish the gut barrier in *Drosophila* larvae and that the lethality associated with gut barrier defects in *Pngl* mutants is primarily caused by non-infectious mechanisms.

## Results

### Loss of *Pngl* is associated with upregulation of immune response-related genes in the midgut

To determine the biological processes that contribute to lethality in *Pngl* mutants, we performed transcriptomic analysis using RNA sequencing (RNA-seq) on third instar larval midguts of *Pngl^–/–^* animals and three control strains: *y w* (*Pngl^+/+^*)*, Pngl^+/–^*, and *Pngl^–/–^; Pngl Dp/+*, which lacks endogenous *Pngl* function, but harbors one copy of a *Pngl* genomic duplication shown to fully rescue the lethality of *Pngl^–/–^* animals^26^. We first identified the genes differentially expressed between *Pngl^–/--^* and each control, as those showing an absolute fold-change of at least 1.5 and a *P-*value less than 0.01. To increase the stringency of our analysis, we focused on those differentially expressed genes that were overlapping among these three pairwise comparisons: (1) *Pngl^–/–^* vs *y w*, (2) *Pngl^–/–^* vs *Pngl^+/–^*, and (3) *Pngl^–/–^* vs *Pngl^–/–^; Pngl Dp/+*. Using this strategy, we found 453 upregulated and 455 downregulated genes in *Pngl^–/–^* larval midguts, when compared to all controls (Fig. 1a and Supplementary Table 1). Using the Database for Annotation, Visualization and Integrated Discovery (DAVID^27,28^, we performed functional gene ontology (GO) analysis on the differentially expressed genes and identified various biological processes significantly altered in *Pngl^–/–^* midguts (Fig. 1b). We found proteasome-mediated processes as the topmost significantly downregulated gene category (Fig. 1b; top panel), in agreement with previous reports on the regulation of proteasomal gene expression by *Pngl* and its homologs^29–33^.

**Figure 1.**
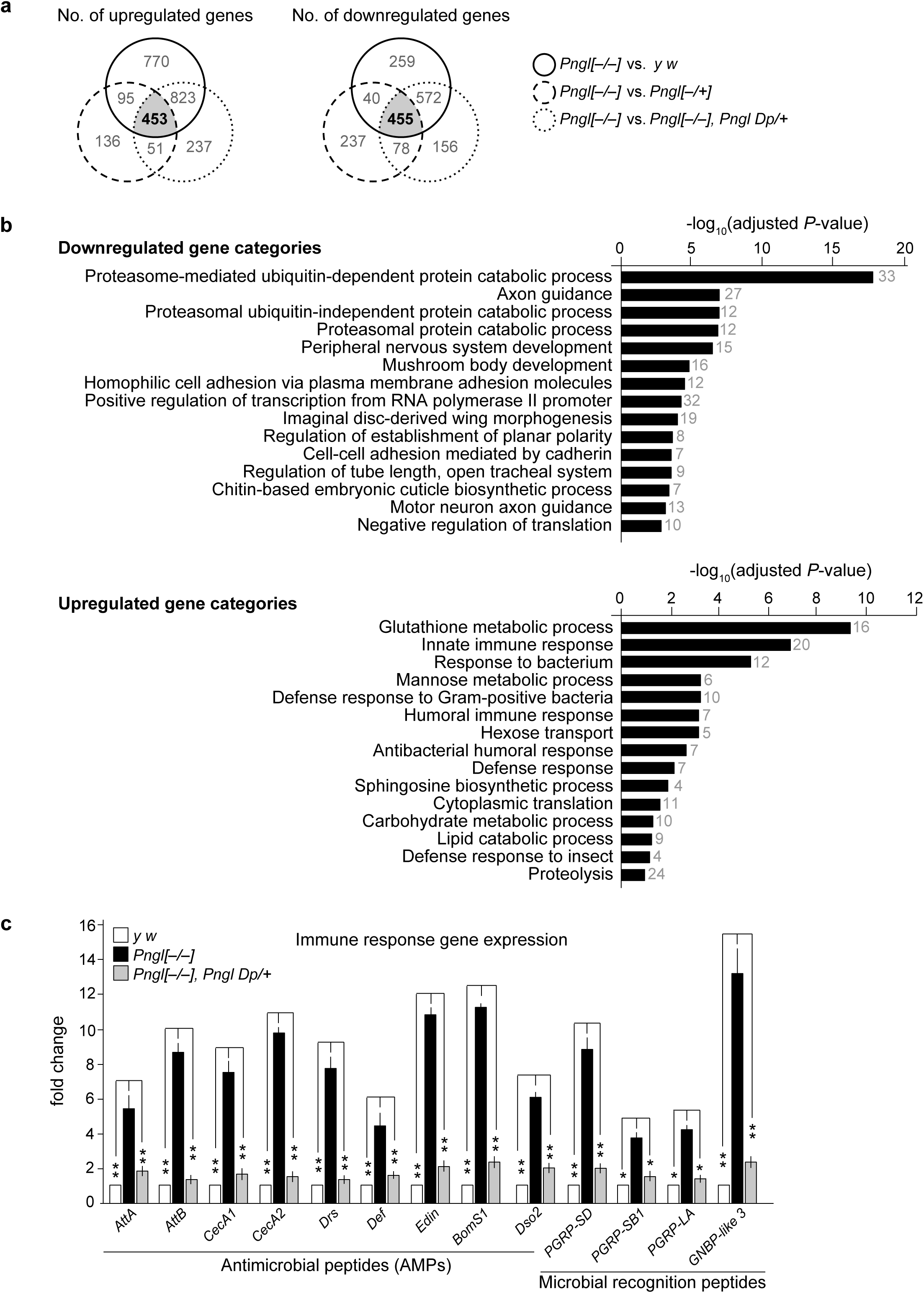
Loss of Pngl is associated with the upregulation of immune response related genes. **a** Venn diagrams showing the overlap of differentially regulated genes in *Pngl^–/–^* with comparison to control (*y w*), *Pngl^+/–^* and *Pngl^–/–^, Pngl Dp/+* in RNA-seq analysis. Number of both upregulated and downregulated genes shown in the Venn diagram were based on >1.5 fold-change. **b** Graph presents DAVID functional GO analysis of biological processes (BP) of downregulated (top), and upregulated (bottom) genes based on their –log10 of adjusted *P*-value. Numbers next to the bars show the number of genes differentially expressed in each category. **c** Graph showing expression of immune response related genes in the indicated genotypes. Values are expressed as fold changes relative to control (*y w*). Error bars represent SD of three replicates. Significance is ascribed as \**P*<0.05 and \*\**P*<0.01 using one-way ANOVA.

Interestingly, a number of significantly upregulated gene categories were related to immune response (Fig. 1b; bottom panel). In *Drosophila,* humoral innate immune response relies on the production of different antimicrobial peptides (AMPs), which can be released systemically or can be synthesized locally in tissues, including the intestinal epithelium, in response to allergens and infection^34,35^. Many of the upregulated genes in the immune response gene categories encode for AMPs and some of them encode for pattern recognition receptor proteins such as peptidoglycan recognition proteins (PGRPs). To confirm the RNA-seq data, we performed qRT-PCR analysis on 13 innate immune response genes from the list and found all of them to be significantly upregulated in *Pngl* mutant midguts (Fig. 1c). Addition of a genomic copy of *Pngl* fully rescued the immune gene expression (Fig. 1c). We conclude that loss of *Pngl* leads to a significant increase in the expression of multiple immune response-related genes in the larval midgut.

Inactivation of NFE2L1 (NFE2 like bZIP transcription factor 1) in NGLY1-deficient *Drosophila* and mouse embryonic fibroblasts (MEFs) leads to impaired proteasomal and mitophagy gene expression^26,30,32^. Moreover, we and others have shown that treatment with an NFE2L2 activator called sulforaphane can rescue these transcriptional defects^26,30,36^. To examine whether NFE2L1 inactivation can explain the innate immune hyperactivation in *Pngl^–/–^* larval midguts, we grew *Pngl^–/–^* larvae on food containing sulforaphane. Despite rescue of proteasomal gene expression by sulforaphane, it did not significantly reduce the expression of innate immunity genes in these animals (Supplementary Fig. 1a and 1b). Moreover, we have reported that increasing the gene dosage of *AMPKα* can significantly rescue the lethality of *Pngl^–/–^* larvae^26^. However, adding an extra copy of *AMPKα* did not rescue the upregulation of most immune response genes in *Pngl^–/–^* midguts and only mildly reduced the expression of some (Supplementary Fig. 2a). Together, these observations indicate that NFE2L1 inactivation and *AMPKα* reduction cannot explain the severe increase in the expression of innate immunity genes in *Pngl^–/–^* larval midguts.

### Enhanced innate immune response in midgut contributes to the lethality of *Pngl* **mutants**

Excessive immune response in the gut can lead to tissue damage and contribute to lethality in adult flies^37,38^. Accordingly, we hypothesized that hyperactive immune response might contribute to the lethality in *Pngl* mutant larvae. Expression of AMPs and other innate immune genes in *Drosophila* is regulated by two signaling pathways, the Toll pathway and the immune deficiency (IMD) pathway^37^. Moreover, in the intestinal epithelium, the forkhead transcription factor Foxo can induce the expression of AMPs in response to starvation, energy deprivation and infection^39,40^. To determine if the increased immune gene expression contributes to the lethality of *Pngl* mutants, we sought to reduce immune activation in these animals by decreasing the gene dosage of *foxo*, *Rel* (encodes Relish, which is the NF-κB transcription factor in IMD pathway) and *Tl* (encodes the Toll receptor). Loss of one copy of *foxo* in *Pngl^–/–^* larvae resulted in a statistically significant reduction in the midgut expression of all 13 immune response genes examined in our experiments, although in most cases the expression levels did not fully return to control levels (Fig. 2a). Loss of one copy of *Rel* and *Tl* also significantly decreased the expression of immune response genes in *Pngl* mutants (Fig. 2a). Notably, we observed a more robust improvement in immune gene expression upon reducing the gene dosage of *foxo* compared to *Rel* and *Tl*, both in terms of the number of genes whose expression was affected and in terms the level of reduction in gene expression (Fig. 2a). *Pngl^–/–^* animals rarely emerge from the pupal case^24,25^. However, reducing the gene dosage of *foxo* rescued the lethality of *Pngl* mutants by 40%, while decreasing the gene dosage of *Rel* and *Tl* rescued the lethality of *Pngl^–/–^* animals by 19% and 21%, respectively (Fig. 2b). Moreover, combined heterozygosity for *foxo* and *Rel* or *Tl* in *Pngl^–/–^* animals did not further increase the degree of lethality rescue achieved by reducing *foxo* gene dosage alone (Fig. 2b). These observations suggest that these genes contribute to *Pngl^–/–^* lethality through a common mechanism, likely the induction of innate immune genes. Of note, adding one genomic copy of *AMPKα* further increased the survival of *Pngl^–/–^; foxo^+/–^* animals (Supplementary Fig. 2b). Together, these observations suggest that *foxo*-mediated hyperactivation of innate immune genes is a major contributor to the lethality in *Pngl* mutants and further suggest that the detrimental effects of innate immune hyperactivation is distinct from the adverse effects of reducing AMPKα signaling in these animals.

**Figure 2.**
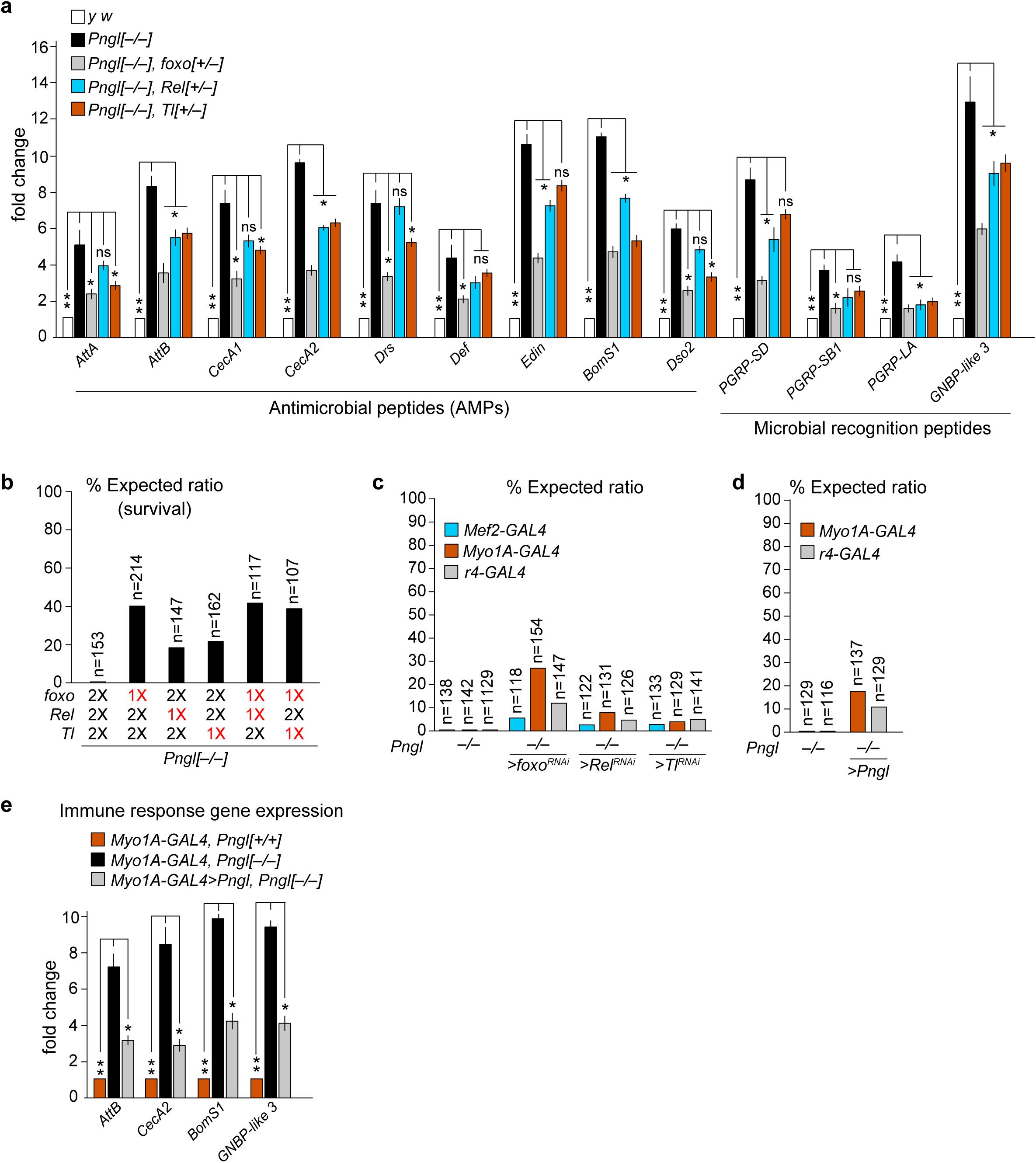
Enhanced innate immune response in midgut contributes to the lethality of *Pngl* mutants. **a** Graph showing immune response genes in the indicated genotypes. Values are expressed as fold changes relative to control (*y w*). Error bars represent SD of three replicates. Significance is ascribed as \**P*<0.05 and \*\**P*<0.01 using one-way ANOVA. ns, not significant. **b** Graph showing % lethality rescue (calculated based on the expected Mendelian ratio) in *Pngl* mutants upon removing one copy (1X) of each immune gene activator. n represents the number of animals scored. **c** Graph showing % lethality rescue in *Pngl* mutants upon tissue-specific knockdown of immune activators using mesoderm-(*Mef2-GAL4*), enterocyte-(*Myo1-GAL4*) and fat body-specific (*r4-GAL4*) drivers. n represents the number of animals scored. **d** Graph showing % lethality rescue in *Pngl* mutants upon enterocyte-and fat body-specific overexpression of *Pngl*. n represents the number of animals scored. **e** Graph showing immune gene expression in the indicated genotypes. Error bars represent SD of three replicates. Significance is ascribed as \**P*<0.05 and \*\**P*<0.01 using one-way ANOVA. ns, not significant.

So far, our gene expression analysis has been performed on whole midguts after removing neuronal and tracheal connections, and our genetic manipulations were performed by using germline mutations. *Drosophila* midgut consists of endoderm-derived epithelium and mesoderm-derived visceral musculature. Therefore, we next performed tissue-specific knockdown experiments to determine the midgut cell type(s) in which hyperactivation of the innate immune response results in *Pngl^–/–^* lethality. We also included fat bodies in our analysis, given their prominent role in systemic release of AMPs^41^. To knockdown *foxo*, *Rel* and *Tl* in the endoderm, mesoderm and fat bodies of *Pngl* mutants, we used *Myo1A-GAL4*, *Mef2-GAL4* and *r4-GAL4* drivers, respectively. Endoderm-specific knockdown of *foxo* (*Myo1A>foxo^RNAi^*) resulted in a lethality rescue of ∼28% in *Pngl* mutants, while fat body specific knockdown (*r4>foxo^RNAi^*) and mesoderm specific knockdown (*Mef2>foxo^RNAi^*) resulted in ∼11% and ∼5% lethality rescue, respectively (Fig. 2c). Knockdown of *Rel* in endoderm (*Myo1A>Rel^RNAi^*), fat body (*r4>Rel^RNAi^*), and mesoderm (*Mef2>Rel^RNAi^*) showed a lethality rescue of ∼8%, 4% and 2%, respectively, in *Pngl* mutants, while knockdown of *Tl* in endoderm (*Myo1A>Tl^RNAi^*) and fat body (*r4>Tl^RNAi^*) showed ∼3% and ∼4% lethality rescue, respectively (Fig. 2c). Taken together, these data suggest that the detrimental effects of immune hyperactivation in *Pngl* mutants primarily results from Foxo-mediated induction of immune response genes in enterocytes and to some extent in the fat body.

Using knockdown and rescue experiments with *Mef2-GAL4* and *how^24B^-GAL4* drivers, we previously showed a major requirement for Pngl in the mesoderm during *Drosophila* development^25,26^. However, knockdown and rescue experiments had suggested that Pngl might not play an important role in the endoderm^25,26^. Given the above observations, we revisited the issue of *Pngl* requirement in endoderm by using the enterocyte-specific driver employed in this study and also examined its requirement in the fat body. Overexpression of *Pngl* using *Myo1A-GAL4* and *r4-GAL4* drivers led to ∼19% and ∼11% rescue of *Pngl^–/–^* lethality, respectively (Fig. 2d). These observations indicate that in addition to its role in mesoderm, Pngl is also required in the endoderm and fat body during *Drosophila* development. Notably, enterocyte-specific overexpression of *Pngl* in *Pngl* mutants led to a significant reduction of innate immune gene expression but did not fully rescue this phenotype (Fig. 2e). These data suggest that although the hyperactive immune response in *Pngl* mutant midguts is primarily due to the loss of *Pngl* in the enterocytes, other cell types and tissues contribute to the induction of immune gene expression in the midgut as well.

### *Pngl* mutant larvae exhibit overactivation of Foxo in their intestine and fat body

Based on the strong rescue of lethality and immune response gene expression achieved by *foxo* heterozygosity in *Pngl* mutants, we examined whether loss of *Pngl* affects Foxo activation. Foxo is negatively regulated by insulin receptor (InR) signaling^42^. Activation of insulin signaling results in phosphorylation of the Akt kinase, which then phosphorylates Foxo to promote its nuclear export^43^. Under conditions of starvation or energy deprivation, decrease in insulin signaling allows the nuclear localization of Foxo and subsequent induction of its target genes^44^. Therefore, we examined the level, subcellular localization and phosphorylation status of Foxo in *Pngl^–/–^* midguts. Immunostaining in the midguts of control and *Pngl* mutant larvae revealed increased nuclear localization of Foxo in *Pngl* mutant midguts (Fig. 3a). In agreement with this observation, western blot analysis showed significantly decreased phospho-Foxo level in *Pngl* mutant midguts (Fig. 3b). These observations suggest increased Foxo activation in midgut upon loss of *Pngl*.

**Figure 3.**
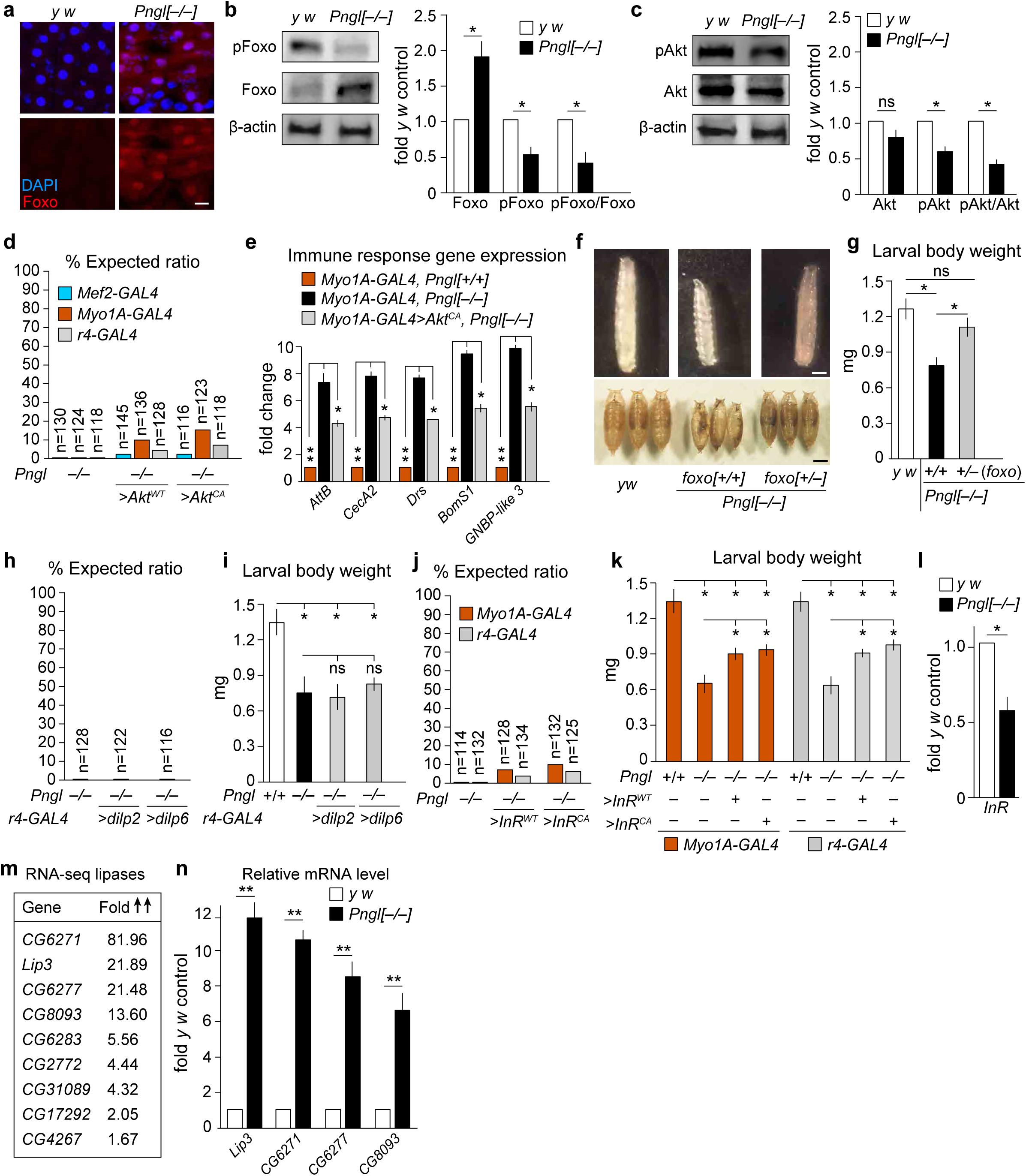
Reduced Akt phosphorylation and enhanced Foxo activation in *Pngl* mutant midguts. **a** Confocal images showing DAPI (blue) and Foxo staining (red) in wild-type and *Pngl* mutant midgut. Scale bar is 50µm. **b** Western blot images and quantification showing the increased Foxo level and decreased phospho-Foxo level in *Pngl* mutant midguts. Error bars represent SD of three replicates. Significance is ascribed as \**P*<0.05 using unpaired student’s t-test. **c** Western blot images and quantification showing decreased Akt phosphorylation in *Pngl* mutants. Error bars represent SD of three replicates. Significance is ascribed as \**P*<0.05 using unpaired student’s t-test. ns, not significant. **d** Graph showing % lethality rescue upon tissue specific overexpression of wild-type (*Akt^WT^*) and constitutively activated (*Akt^CA^*) form of *Akt* in *Pngl* mutants. n represents the number of animals scored. **e** Graph showing expression of immune genes in the indicated genotypes. Error bars represent SD of three replicates. Significance is ascribed as \**P*<0.05 and \*\**P*<0.01 using one-way ANOVA. **f** Images of larvae and pupae of the indicated genotypes. **g** Graph showing quantification of larval body weight of the indicated genotypes. Error bars represent SD of three replicates. Significance is ascribed as \**P*<0.05 using one-way ANOVA. **h** Graph showing % lethality rescue upon tissue-specific overexpression of *Drosophila* insulin-like peptides (*dilp2* and *dilp6*) in *Pngl* mutants. n indicate the number of animals scored. **i** Graph showing quantification of body weight of indicated genotypes. Error bars represent SD of three replicates. Significance is ascribed as \**P*<0.05 using one-way ANOVA. **j** Graph showing % lethality rescue upon tissue-specific overexpression of wild-type (*InR^WT^*) and constitutively activated (*InR^CA^*) forms of *InR* in *Pngl* mutants. n indicate the number of animals scored. **k** Graph showing larval body weight upon tissue-specific overexpression of *InR^WT^* and *InR^CA^* in *Pngl* mutants (n>20 for each condition). Error bars represent SD of three replicates. Significance is ascribed as \**P*<0.05 using one-way ANOVA. **l** Graph showing the expression of *InR* in the indicated genotypes. Error bars represent SD of three replicates. Significance is ascribed as \**P*<0.05 using unpaired student’s t-test. **m** List of genes in the “Lipid catabolic process” from the larval midgut RNA-seq, along with their fold increase in expression level in *Pngl^–/–^* compared to *y w* control. **n** Graph showing the increased expression of lipases in *Pngl* mutant midgut. Error bars represent SD of three replicates. Significance is ascribed as \*\**P*<0.01 using unpaired student’s t-test.

We next examined the phosphorylation status of Akt in *Pngl* mutants. Western blot analysis showed a significant decrease in phospho-Akt levels in *Pngl* mutants (Fig. 3c), suggesting reduced Akt phosphorylation contributes to overactivation of Foxo in these animals. In support of this notion, we observed a lethality rescue of ∼11% and ∼17% by enterocyte-specific overexpression of wild-type Akt (*Akt^WT^*) and constitutively active Akt (*Akt^CA^*), respectively, in *Pngl* mutants (Fig. 3d). Furthermore, overexpression of *Akt^CA^* in enterocytes also led to a significant reduction in innate immune gene expression in *Pngl* mutants (Fig. 3e). Together, these data suggest that reduced Akt activation in *Pngl* mutant midguts contributes to Foxo overactivation and subsequently increased immune gene expression and lethality.

As mentioned before, Akt is regulated by insulin signaling, and Foxo is a key transcriptional effector of the insulin signaling pathway^43^. Moreover, in agreement with a previous report^45^, we observed a small body size phenotype (based on body weight) in *Pngl* mutant third instar larvae (Fig. 3f, g). This phenotype, which is a signature of decreased insulin signaling in flies^46^, was also rescued by decreasing the gene dosage of *foxo* (Fig. 3f, g). Accordingly, our data suggest that reduced insulin signaling in the midgut might contribute to the lethality of *Pngl* mutants. To test this notion, we examined the effects of overexpressing insulin ligands or receptor on *Pngl^–/–^* phenotypes. The *Drosophila* insulin-like peptide 6 (Dilp6; official name Ilp6) is primarily expressed in third instar larval fat body and is essential for animal growth^47^. However, overexpression of *dilp6* in the fat body did not rescue the lethality of *Pngl* mutants (Fig. 3h). Analysis of the *Drosophila* Dilps by NetNGlyc - 1.0 sever (https://services.healthtech.dtu.dk/service.php?NetNGlyc-1.048) indicates that none of the *Drosophila* Dilps harbor predicted *N*-glycosylation sites except for Dilp6, which has a predicted *N*-glycosylation sequon at ^85^**N**VT^87^. We have recently reported that the enzymatic activity of Pngl and its mammalian homolog is required for deglycosylation of misfolded Dpp/BMP4 ligands and secretion of mature Dpp/BMP4 is some cell types^49^. Therefore, to account for a potential effect of loss of *Pngl* on the secretion of Dilp6, we also overexpressed Dilp2 in the fat bodies of *Pngl* mutants. Dilp2 overexpression by *r4-GAL4* did not rescue the *Pngl^–/–^* lethality (Fig. 3h). Overexpression of *dipl2* and *dilp6* did not improve the small body size phenotype either (Fig. 3i). Next, we overexpressed wild-type InR (*InR^WT^*) and constitutively activate InR (*InR^CA^*) in *Pngl* mutants using *Myo1A-GAL4* and *r4-GAL4* drivers. We observed ∼8% and ∼10% rescue of the lethality upon overexpression of *InR^WT^* and *InR^CA^* in the enterocytes of *Pngl* mutants, respectively (Fig. 3j). A lower degree of rescue was observed upon *InR^WT^* and *InR^CA^* overexpression in the fat body (Fig. 3j). Overexpression of *InR^WT^* and *InR^CA^* in enterocytes and fat body also exhibited a partial rescue of body size in *Pngl* mutants (Fig. 3k). Moreover, we observed a significant decrease in the expression of *InR* in the midgut of *Pngl* mutants (Fig. 3l). These observations suggest that reduced InR signaling in the midgut and potentially fat body contributes to the lethality of *Pngl* mutants.

Reduced InR signaling can result from starvation^50^. Therefore, we examined whether *Pngl^–/–^* larvae show any evidence for starvation. One of the hallmarks of starvation in *Drosophila* larvae is an increase in lipid catabolism. Importantly, one of the upregulated gene categories in the RNA-seq analysis was related to lipid catabolic process (Fig. 1b; bottom panel and Fig. 3m), which was confirmed by qRT-PCR experiments showing a significant increase in the expression of several lipases from this list (*Lip3*, *CG6271*, *CG8093* and *CG6277*) (Fig. 3n). These observations suggest that starvation contributes to reduced InR signaling and Foxo activation in *Pngl* mutant larvae.

### Loss of *Pngl* is associated with gut barrier defects, which contribute to the lethality in *Pngl* mutants

Gut barrier defects can lead to activation of the intestinal innate immune response^2,3^. Therefore, we examined whether *Pngl^–/–^* larvae exhibited any gut barrier dysfunction. We fed the control and *Pngl*-mutant larvae on food containing FITC-labeled high molecular weight (500-kDa) dextran and imaged their midguts to visualize the dextran. In the midgut of control larvae, the FITC signal was restricted to the central parts of the lumen area, but the *Pngl^–/–^* midguts failed to retain the FITC signal in the lumen area and showed strong FITC signal filling the spaces among neighboring gut epithelial cells (Fig. 4a and 4b). Around 55% of the examined *Pngl*-mutant third instar larvae (but none of the WT) showed gut barrier defect, and providing one genomic copy of *Pngl* fully rescued the phenotype (Fig. 4c). As shown in Supplementary Fig. 2a, increasing the gene dosage of *AMPKα* reduced the expression of some innate immunity genes in *Pngl^–/–^* larval midguts. However, adding one copy of *AMPKα* did not rescue the gut barrier defects in *Pngl^–/–^* larvae (Supplementary Fig. 2c). These observations indicate that Pngl is required for normal gut barrier formation in *Drosophila* larvae independently of the AMPKα defects previously reported in these animals.

**Figure 4.**
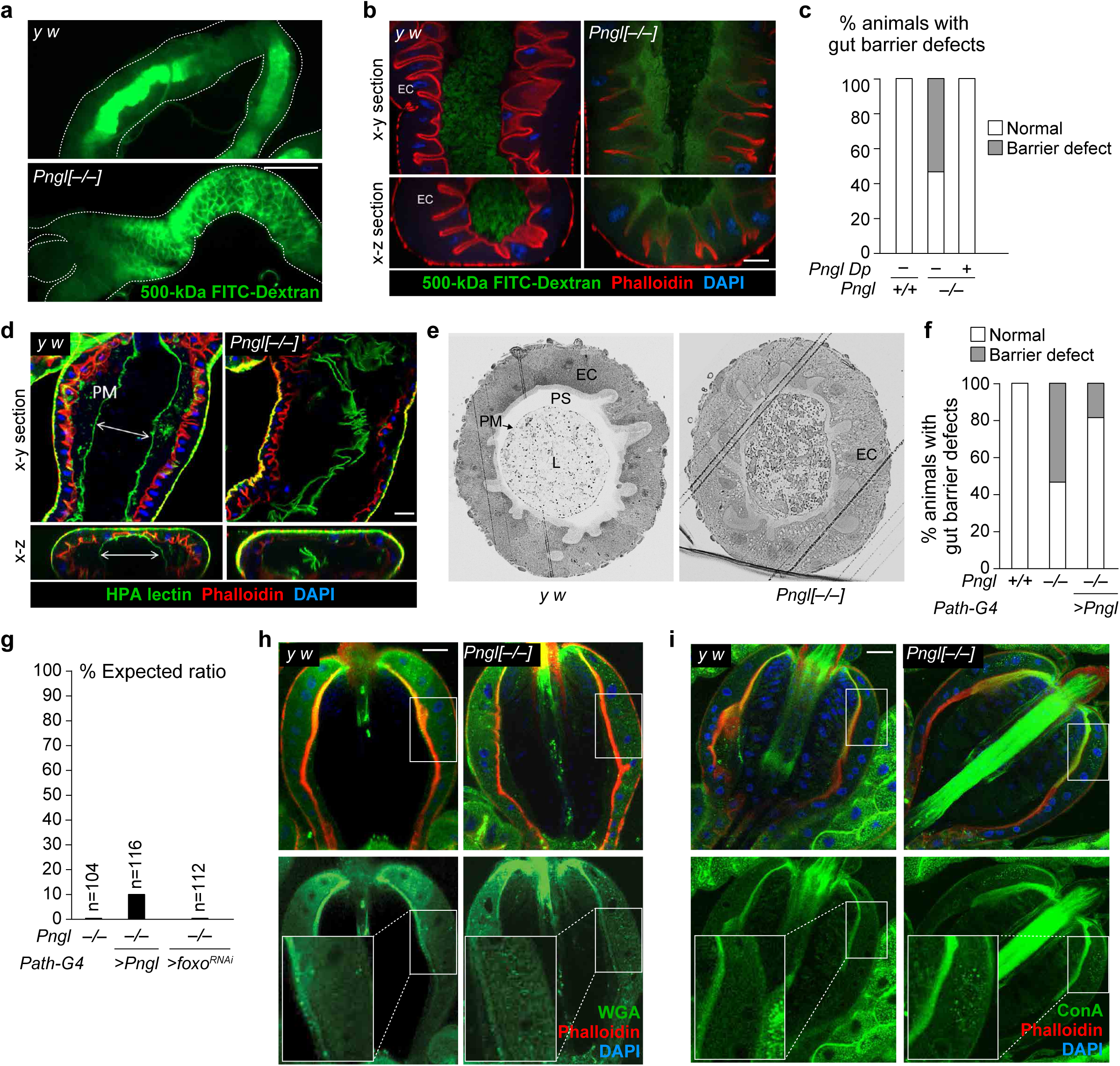
Loss of Pngl is associated with gut barrier defect which contributes to the lethality in *Pngl* mutants. **a** Low magnification fluorescent images of wild-type and *Pngl* mutant midgut upon FITC-labeled dextran (Green) feeding. **b** Confocal images showing phalloidin (red) and DAPI (blue) staining in wild-type and *Pngl* mutant midgut upon FITC-labeled dextran (Green) feeding. Scale bar is 50 µm. **c** Graph showing the quantification of gut barrier defect phenotype in the indicated genotypes (n=60 for each genotype). **d** Confocal image showing phalloidin (red), DAPI (blue) and HPA-lectin staining (Green) for peritrophic matrix in wild-type and *Pngl* mutant midgut. Scale bar is 50 µm. **e** Light microscopic images of cross-section of wild-type and *Pngl* mutant midguts. Lumen (L), Peritrophic matrix (PM), Peritrophic space (PS), Enterocytes (EC). **f** Graph showing quantification of gut barrier defect phenotype in the indicated genotypes (n=60 for each genotype. **g** Graph showing lethality rescue in *Pngl* mutants upon *foxo* knockdown and *Pngl* overexpression in their PM secretory region. n indicates the number of animals scored. **h** Confocal images showing Phalloidin (red), DAPI (blue) and WGA lectin staining (Green) in the proventriculus region of wild-type and *Pngl^–/–^* third instar larvae (n=4 for each genotype). Insets show close-up of PM-secreting cells. Scale bar is 50 µm. **i** Confocal images showing Phalloidin (red), DAPI (blue) and ConA lectin staining (Green) in the proventriculus region of wild-type and *Pngl^–/–^* third instar larvae (n=5 for each genotype). Brackets mark the PR cells. Scale bar is 50 µm.

Given the abnormality observed in the gut barrier function of *Pngl^–/–^* larvae, we sought to examine the integrity of peritrophic matrix (PM) in these larvae. To this end, we marked the PM in *Pngl^–/–^* and control larvae with Helix pomatia agglutinin (HPA) lectin, which selectively binds to α-*N*-acetylgalactosamine residues and is specific for *O-* glycans^51,52^, and performed confocal imaging. In control larvae, PM separated a central luminal area from a peripheral “peritrophic” space adjacent to the apical surface of the midgut epithelium (Fig. 4d), an arrangement which is thought to ensure that abrasive food particles and microorganisms pass through the gut without contacting the epithelial cells^13^. However, although PM can clearly be observed in the midgut of *Pngl^–/–^* larvae, it is highly disorganized and appears to be collapsed on itself (Fig. 4d). This is accompanied by epithelial irregularities, as evidenced by defects in apical phalloidin staining and detachment of some epithelial cells (Fig. 4d). In addition, analysis of midgut sections by light microscopy showed the lumen area lined with intact PM and a peritrophic space between enterocytes and PM in control animals. In contrast, *Pngl* mutant midguts displayed a dense lumen content (potentially due to the gut clearance defect previously reported in these animals^26^), disorganized PM, and loss of peritrophic space (Fig. 4e). These observations indicate impaired gut barrier function in *Pngl^–/–^* larvae and abnormalities in PM.

To determine if loss of *Pngl* in PR cells contributes to the gut barrier defects observed in *Pngl^–/–^* larvae, we overexpressed *Pngl* in PR cells of *Pngl* mutants and fed the larvae with 500-kDa FITC-dextran. We found that the penetrance of gut barrier phenotype in *Pngl* mutant larvae was reduced from 55% to ∼20% upon PR-specific overexpression of *Pngl* (Fig. 4f). This was accompanied by a lethality rescue of ∼11% (Fig. 4g). Importantly, *foxo* knockdown in PR cells did not lead to any rescue of lethality (Fig. 4g). Together, these data suggest that Pngl is required in PR cells to ensure the integrity of the PM and gut barrier and to promote the survival of *Drosophila* larvae.

Loss of NGLY1 and its homologs affects the degradation of some misfolded *N*-glycoproteins and lead to their cytosolic accumulation^19,53–55^. Moreover, we have previously reported that upon loss of *NGLY1*, misfolded BMP4 molecules fail to be degylcosylated and retrotranslocated from the ER to the cytosol, leading to accumulation of misfolded BMP4 in the ER and a failure in the secretion of properly folded BMP4^49^. To assess whether there is an accumulation of *N*-glycoproteins in the PR cells of *Pngl* mutant larvae, we stained the proventricular regions of these animals and control third instar larvae with two lectins: wheat germ agglutinin (WGA), which recognizes *N*-acetylglucosamine in *O*-glycans, chitins, and *N*-glycans, and concanavalin A (Con A), which primarily binds high-mannose *N*-glycans^15,56^. Notably, *Pngl* mutants show an accumulation of WGA^+^ and ConA^+^ intracellular puncta in the PR cells, suggesting that loss of *Pngl* might affect the trafficking and/or secretion of some *N*-glycoproteins (Fig. 4h and 4i). These observations provide a potential mechanism for the PM defects observed in *Pngl^–/–^* larvae.

### Germ-free rearing does not rescue the gut barrier defect and lethality of *Pngl* **mutants**

The gut barrier defects and upregulation of multiple microbial recognition peptides in *Pngl^–/–^* larvae (Fig. 4 and Fig. 1c) suggest that the gut microbiota might contribute to some of the *Pngl^–/–^* phenotypes. Moreover, altered gut microbiota can precede the intestinal barrier defects in aging adult *Drosophila*^57^. To directly examine whether gut microbiota play any roles in *Pngl-*mutant phenotypes, we generated germ-free *Pngl^–/–^* and control larvae. Germ-free rearing of *Pngl* mutants did not lead to any improvement in their gut barrier phenotype (Fig. 5a), suggesting that the PM defects caused by loss of *Pngl* are not due to altered gut microbiota.

**Figure 5.**
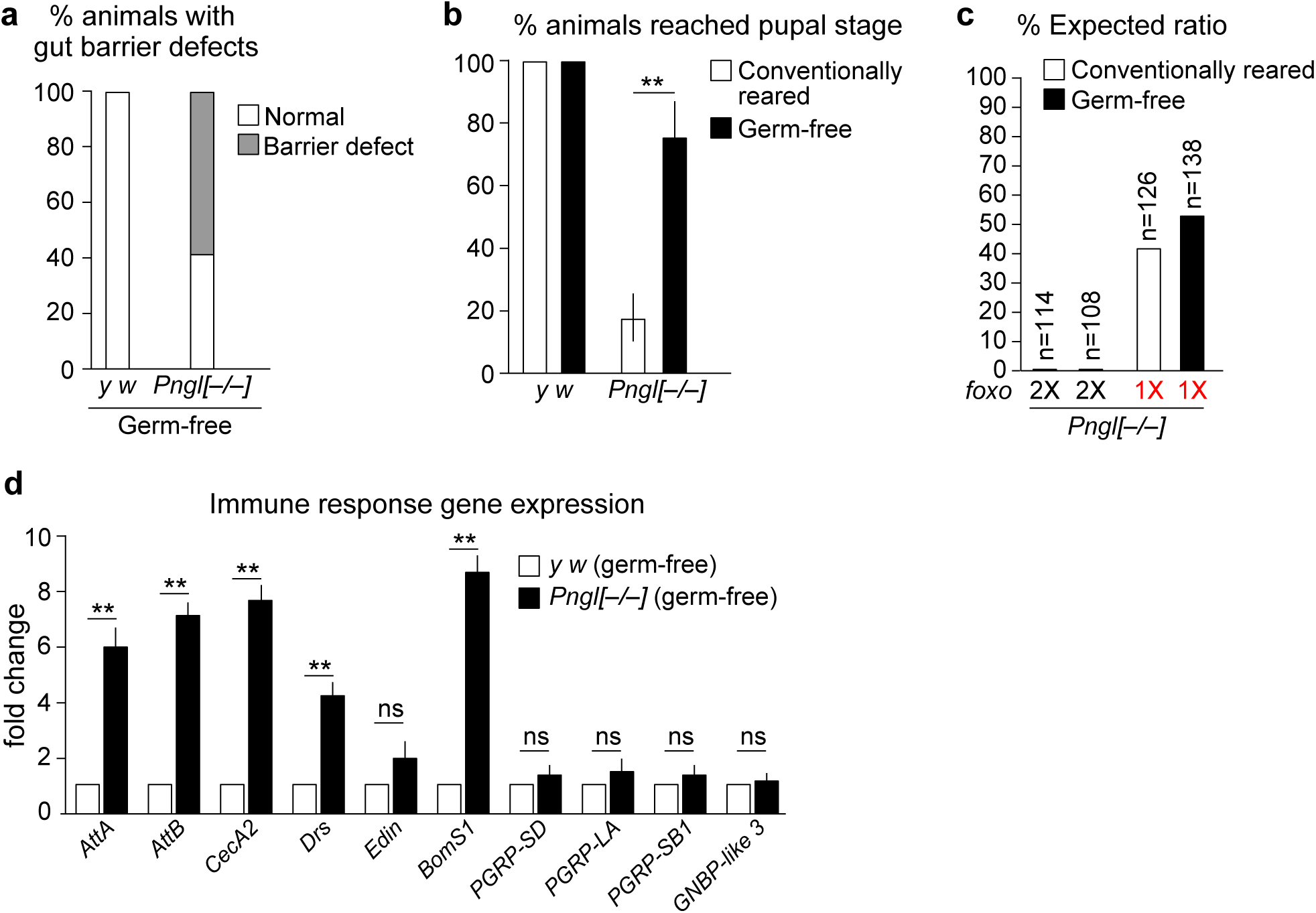
Gut microbial abundance does not explain gut barrier defects and lethality in *Pngl* mutants. **a** Graph showing quantification of gut barrier defect phenotype in germ-free wild-type and *Pngl* mutants (n=60 for each condition). **b** Graph showing the percentage of germ-free and conventionally reared wild-type and *Pngl^–/–^* animals that reached the pupal stage (n= 90 for each condition). Error bars represent SD of three replicates. Significance is ascribed as \*\**P*<0.01 using unpaired student’s t-test. **c** Graph showing % lethality rescue in germ-free and conventionally reared *Pngl^–/–^* and *Pngl^–/–^; foxo^+/–^*. n indicates the number of animals scored. **d** Graph showing immune gene expression in germ-free wild-type and *Pngl* mutants. Error bars represent SD of three replicates. Significance is ascribed as \*\**P*<0.01 using unpaired student’s t-test. ns, not significant.

The germ-free *Pngl^–/–^* larvae showed a significant increase in the percentage of larvae which reach the pupal stage compared to conventionally reared *Pngl^–/–^* larvae (Fig. 5b), suggesting a role for gut dysbiosis in the developmental delay of *Pngl* mutants. However, germ-free rearing did not lead to any rescue in the lethality of *Pngl* mutants (Fig. 5c). Of note, germ-free rearing of *Pngl^–/–^; foxo^+/–^* larvae resulted in further improvement in their survival compared to conventionally reared *Pngl^–/–^; foxo^+/–^* animals (Fig. 5c, also compare to Fig. 2b). Quantitative RT-PCR showed that expression of the genes involved in the recognition of microorganisms is not statistically different between germ-free *Pngl^–/–^* and germ-free *y w* larval midguts (Fig. 5d). However, the germ-free *Pngl* mutants still exhibit a significant increase in the expression of most of AMPs compared to germ-free control larvae (Fig. 5d). Together, these data suggest that altered gut microbiota is not the major inducer of AMP gene expression in *Pngl* mutants, and that the contribution of gut microbiome to the lethality in *Pngl* mutants is redundant to that of Foxo hyperactivation.

### Gut barrier defects lead to starvation and contribute to Foxo-dependent innate immune gene induction and lethality in *Pngl* mutants

So far, our data suggest that gut dysbiosis cannot explain the detrimental effects of gut barrier defects on the survival of *Pngl^–/–^* larvae. To gain mechanistic insight into the role of gut barrier defects in the lethality of *Pngl^–/–^* larvae and to separate the effects of gut barrier defect from the cell-autonomous effects of loss of *Pngl* in the midgut and fat body, we induced gut barrier defects in control (*Pngl^+/+^*) animals by feeding them with polyoxin D (Poly D), a chitin synthase inhibitor that disrupts PM formation in insects^58,59^. Poly D feeding for 48 hours resulted in a 43%-penetrant gut barrier phenotype, ∼19% lethality, and increased expression of immune genes in *y w* animals (Fig. 6a–c), confirming that this strategy can impair PM formation in *Drosophila* larvae.

**Figure 6.**
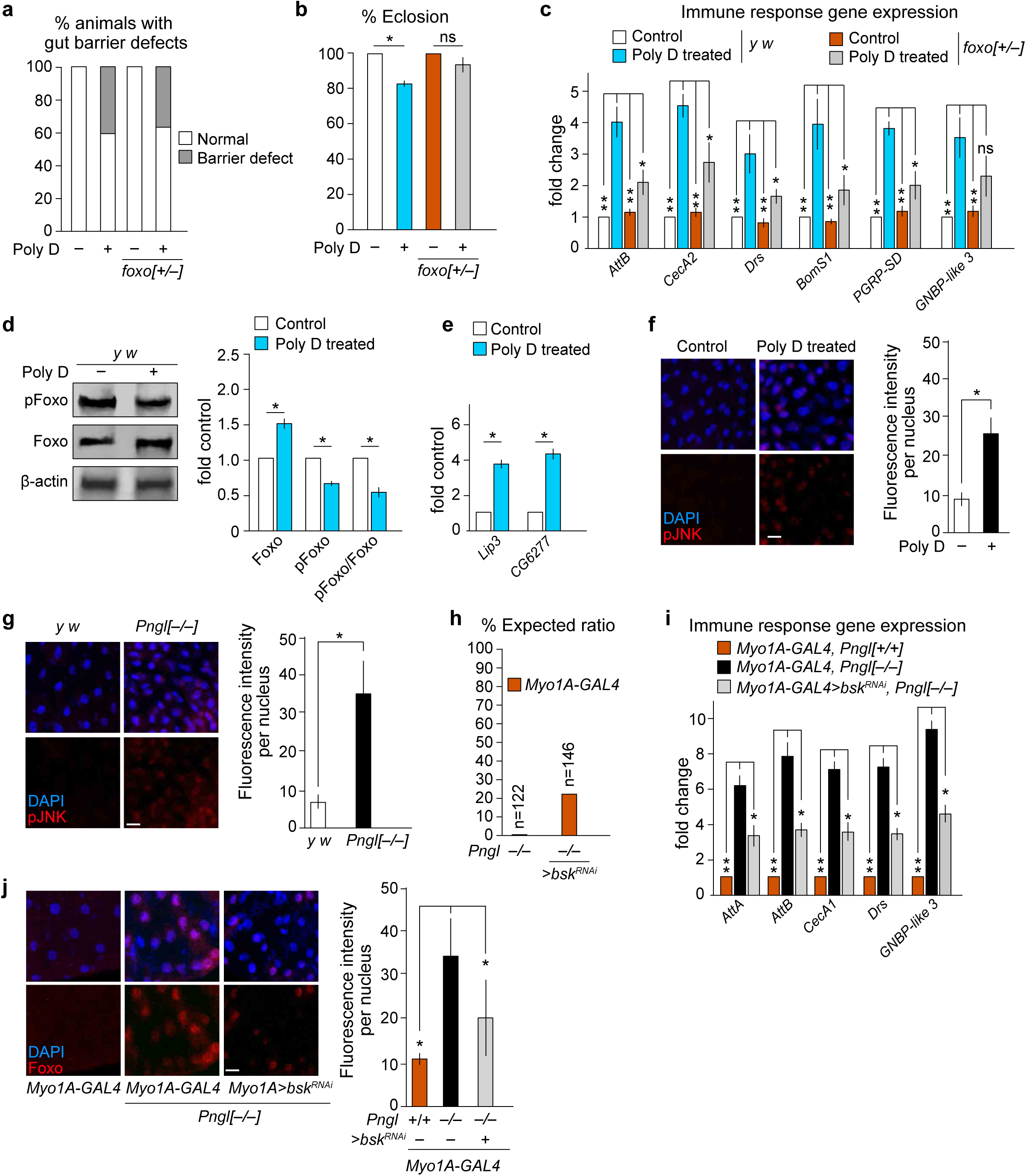
Gut barrier defects induce starvation and Foxo-dependent induction of innate immune genes and lethality. **a** Graph showing the quantification of gut barrier defect in control and polyoxin-D (Poly D) fed wild-type and *foxo^+/–^* mutants (n=60 per condition). **b** Graph showing % adult eclosion in Poly D fed wild-type and *foxo^+/–^* mutants (n=60 per condition). Error bars represent SD of three replicates. Significance is ascribed as \**P*<0.05 using unpaired student’s t-test. ns, not significant. **c** Graph showing immune response gene expression in control and Poly D fed wild-type and *foxo^+/–^* mutant midguts. Error bars represent SD of three replicates. Significance is ascribed as \**P*<0.05 and \*\**P*<0.01 using two-way ANOVA. ns, not significant. **d** Western blot images and quantification graph showing increased activation of Foxo in Poly D fed wild-type midguts. Error bars represent SD of three replicates. Significance is ascribed as \**P*<0.05 using unpaired student’s t-test. **e** Graph showing expression of lipases (*Lip3* and *CG6277*) in Poly D fed wild-type midguts. Error bars represent SD of three replicates. Significance is ascribed as \**P*<0.05 using unpaired student’s t-test. **f** Confocal images showing DAPI (blue) and pJNK staining (red) and quantification in control and Poly D fed wild-type midguts. Scale bar is 50 µm. Error bars represent SD of three replicates. Significance is ascribed as \**P*<0.05 using unpaired student’s t-test. **g** Confocal images showing DAPI (blue) and pJNK staining (red) and quantification in wild-type and *Pngl* mutant midguts. Scale bar is 50 µm. Error bars represent SD of three replicates. Significance is ascribed as \**P*<0.05 using unpaired student’s t-test. **h** Graph showing % lethality rescue upon enterocyte-specific knockdown of *Drosophila* JNK (*bsk*) in *Pngl* mutants. n indicates number of animals scored. **i** Graph showing innate immune gene expression in the indicated genotypes. Error bars represent SD of three replicates. Significance is ascribed as \**P*<0.05 and \*\**P*<0.01 using one-way ANOVA. **j** Confocal images showing DAPI (blue) and Foxo staining (red) and quantification of Foxo nuclear localization in the indicated genotypes. Scale bar is 50 µm. Error bars represent SD of three replicates. Significance is ascribed as \**P*<0.05 using one-way ANOVA.

Poly D fed *foxo^+/–^* larvae showed a 38%-penetrant gut barrier phenotype, indicating that as expected, loss of one copy of *foxo* does not affect the ability of Poly D to impair gut barrier integrity (Fig. 6a). However, removing one copy of *foxo* rescued the lethality caused Poly D feeding and significantly reduced the expression of most immune genes induced by the chemical (Fig. 6b and 6c). Further, western blot analysis revealed a significant reduction in the relative levels of phospho-Foxo in midguts of Poly D-fed larvae, suggesting Foxo activation (Fig. 6d). These data strongly suggest that the detrimental effects of impaired gut barrier on *Drosophila* larval development primarily result from Foxo-mediated induction of innate immunity in the midgut.

In dextran feeding assays (Fig. 4) and bromophenol blue feeding assays^25^, loss of *Pngl* did not appear to affect the amount of food intake in *Drosophila* larvae. Moreover, quantification of the number of mandibular movements per second suggests a comparable feeding behavior in control and *Pngl*-mutant larvae (Supplementary Fig. 3). Therefore, the starvation signature observed in these animals is not likely to be due to reduced feeding. Importantly, the Poly D-fed *y w* larvae showed a statistically significant increase in the expression of lipases in the midgut, suggesting a starvation condition (Fig. 6e). Together, these observations indicate that gut barrier defects can lead to a starvation-like condition in *Drosophila* larvae and suggest that Foxo likely operates downstream of this starvation-like phenotype to mediate the lethality associated with gut barrier defects.

Overexpression of InR or Akt in the midgut was less effective in rescuing the *Pngl^–/–^* lethality compared to *foxo* knockdown by using the same driver (compare Fig. 3d and 3j to Fig. 2c). Therefore, in addition to reduced InR signaling, other pathways are likely to contribute to Foxo activation and lethality in *Pngl^–/–^* midguts. The Jun-N-terminal Kinase (JNK) pathway is a potential candidate for these effects, as it is activated in epithelial tissues by a variety of extrinsic and intrinsic stressors and can induce the nuclear localization and activation of Foxo^60,61^. Indeed, feeding Poly D to wild-type larvae led to the activation of JNK signaling in larval midgut, as evidenced by the accumulation of phospho-JNK (pJNK) in their midgut epithelial cells (Fig. 6f). Similarly, *Pngl^–/–^* larvae accumulated pJNK in their midgut epithelial cells (Fig. 6g), supporting the notion that gut barrier defects induce JNK signaling in *Pngl^–/–^* midguts.

To evaluate the functional significance of JNK activation in *Pngl^–/–^* larvae, we overexpressed double-stranded RNA for the *Drosophila* JNK homolog *basket* (*bsk*) in the enterocytes of *Pngl^–/–^* larvae. We observed a lethality rescue of 22% upon *bsk* knockdown in midgut epithelial cells of *Pngl^–/–^* larvae (Fig. 6h). This was accompanied by a significant decrease in immune gene expression and a significant decrease in Foxo nuclear localization in midguts (Fig. 6i and 6j). Together, these observations suggest that gut barrier defects lead to increased JNK signaling in *Pngl^–/–^* midgut epithelial cells, which itself contributes to Foxo activation and lethality in these animals.

### *Pngl* mutants exhibit an increase in lipid catabolism and isocaloric dietary lipid supplementation partially rescues the lethality in these animals

Starvation can lead to lipid mobilization^50,62^. In line with the significant increase in the expression of multiple lipase genes in *Pngl^–/–^* midguts (Fig. 3l and 3m), we observed a significant reduction in the lipid storage in fat body and midgut of *Pngl* mutants (Fig. 7a). Triacylglycerol (TAG) is the main energy reservoir in *Drosophila*^63^. In accordance with the dramatic increase in the expression of multiple lipase genes and reduction of fat storage in *Pngl^–/–^* tissues, we found a significant decrease in TAG level in hemolymph and midgut of *Pngl* mutant throughout the third instar larval period (Fig. 7b). This was accompanied by a significant increase in free fatty acid levels in hemolymph of early third instar *Pngl* mutants (Fig. 7c, 72 hr). By mid-third instar larval stage, free fatty acid levels return to normal (Fig. 7c, 108 hr), potentially suggesting the depletion of energy reserve as the development of *Pngl^–/–^* larvae proceeds. Loss of one copy of *foxo* improved all of these phenotypes (Fig. 7a–c). Together, these data indicate that *Pngl^–/–^* larvae experience a significant degree of starvation associated with a Foxo-dependent increase in lipid catabolism.

**Figure 7.**
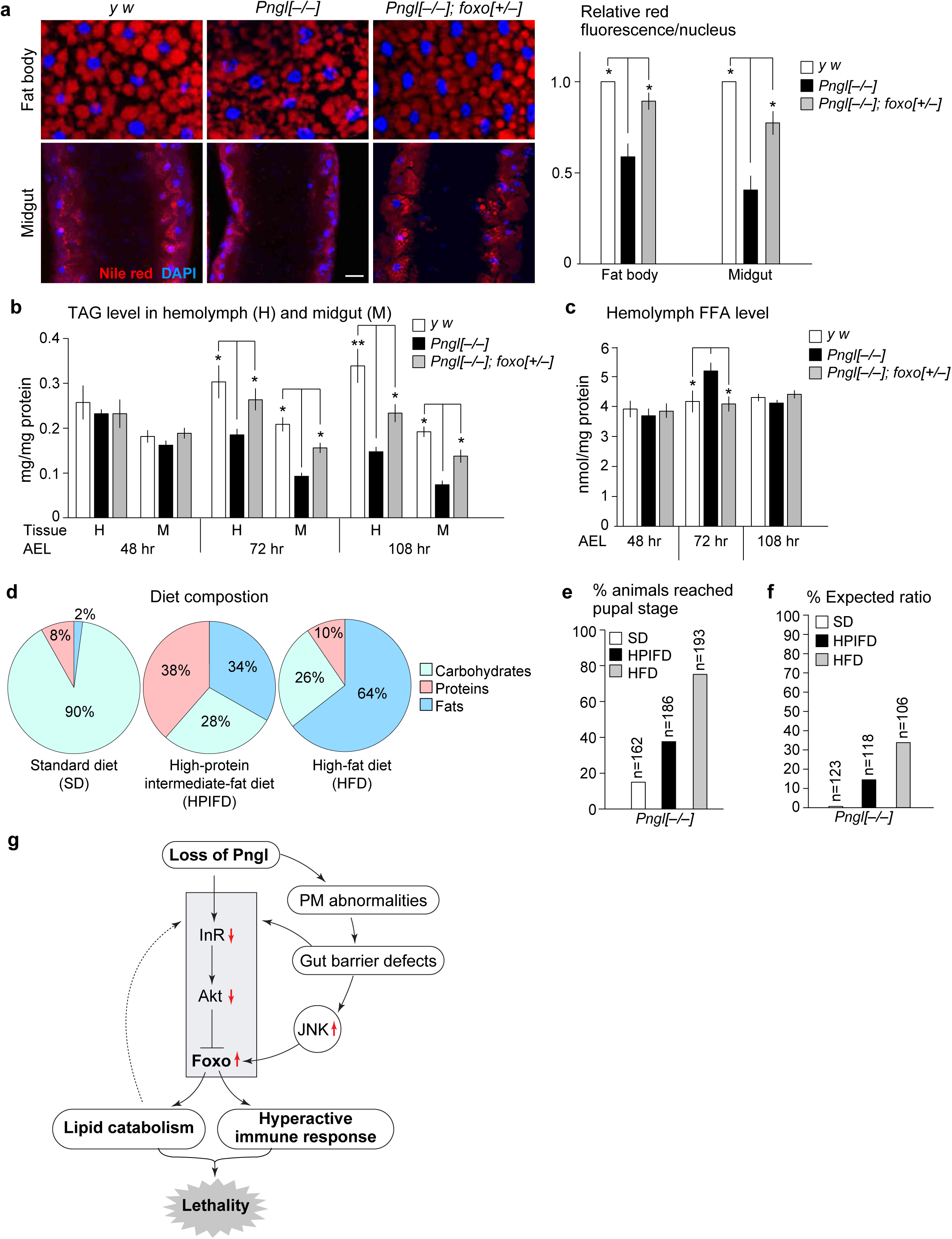
Pngl mutant exhibit increased lipid catabolism and supplementation on dietary lipid partially rescues lethality in *Pngl* mutants. **a** Images showing Nile red (red) and DAPI (blue) staining in fat body (top) and midgut (bottom) of the indicated genotypes and their quantification. Scale bar is 50 µm. Error bars represent SD of three replicates. Significance is ascribed as \**P*<0.05 using one-way ANOVA. b Triacylglycerol (TAG) level in hemolymph and midgut of indicated genotypes. Error bars represent SD of three replicates. Significance is ascribed as \**P*<0.05 and \*\**P*<0.01 using one-way ANOVA. c Free fatty acid (FFA) level in the hemolymph of indicated genotypes. Error bars represent SD of three replicates. Significance is ascribed as \**P*<0.05 using one-way ANOVA. d Pie charts showing % carbohydrates, proteins and fats in different diet compositions. e Graph showing % animals that reach to pupal stage upon feeding on the indicated diets. n indicates the number of animals scored. f Graph showing lethality rescue in *Pngl* mutants upon feeding on the indicated diets. n indicates the number of animals scored. g Schematic model showing that loss of *Pngl* results in Foxo overactivation and subsequent innate immune gene expression and lipid catabolism, leading to lethality.

In the standard fly food used in our experiments, only 2% of the total energy contents is provided by lipids. The depletion of lipid storage in *Pngl^–/–^* midgut and fat body prompted us to examine whether increasing the fat content of the food can promote the survival of *Pngl^–/–^* larvae. To this end, we used two additional isocaloric food compositions with different fat contents in our experiments: a “high-protein intermediate-fat diet” (HPIFD) and a “high-fat diet” HFD (Fig. 7d and Supplementary Table 2). On our standard diet (SD), 14.8% of *Pngl^–/–^* larvae reached the pupal stage (Fig. 7e). However, when grown on isocaloric HPIFD and HFD, 38.3% and 74.1% of *Pngl^–/–^* larvae reached the pupal stage, respectively (Fig. 7e). Further analysis indicated that the developmental delay of *Pngl^–/–^* larvae was partially rescued by HPIFD and fully rescued by HFD (Supplementary Fig. 4A). Remarkably, HPIFD and HFD also led to *Pngl^–/–^* lethality rescues of 14.5% and ∼36%, respectively (Fig. 7f). Immunostaining indicated that HFD significantly reduced the Foxo nuclear localization in midgut and fat body of *Pngl* mutants (Supplementary Fig. 4B), and partially rescued innate immune gene overexpression in *Pngl* mutants (Supplementary Fig. 4c). These data underscore the contribution of metabolic abnormalities and energy depletion to the developmental delay and lethality in *Pngl* mutants.

## Discussion

We previously reported that Pngl is required in the midgut visceral mesoderm for the regulation of Dpp and AMP kinase signaling in *Drosophila*^25,26,49^. However, impairment of these two pathways only partially explained the lethality of *Pngl^–/–^* animals. Here, we report that *Pngl* plays critical roles in several other cell types (PR cells, enterocytes, and fat body cells). Loss of *Pngl* leads to gut barrier defects, causing suppression of InR signaling and activation of JNK signaling in enterocytes. These alterations result in overactivation of Foxo in enterocytes and fat body, which in turn leads to hyperactivation of intestinal innate immune genes and increased lipid catabolism, ultimately causing lethality (Fig. 7g). *Pngl* mutants show significant developmental delay, with less than 1% of the animals reaching the end of the pupal stage and eclosing as an adult^24,25^. Intriguingly, growing *Pngl* mutants on an isocaloric high-fat diet fully rescued their developmental delay and allowed 33% of the animals to reach adulthood, strongly suggesting that depletion of energy depot during the larval and pupal development plays a critical role in the lethality of *Pngl* mutants. We note that a high-fat diet also suppressed Foxo overactivation in the fat body and midgut of *Pngl^–/–^* larvae. This observation suggests that by worsening the starvation status, enhanced lipid catabolism engages in a vicious cycle with Foxo overactivation and negatively impacts the survival of *Pngl^–/–^* animals.

Our data provide strong evidence that hyperactivation of innate immune genes contributes to lethality in *Pngl* mutants. First, decreasing the gene dosage of *foxo*, *Rel* and *Tl*, which are the known regulators of innate immune gene expression, partially rescued the lethality in *Pngl* mutants. In addition, in all of our other genetic and diet-induced rescue experiments, rescue of lethality was accompanied by reduced immune gene expression. It has recently been reported that overexpression of AMPs results in cytotoxicity and cell death in aging *Drosophila*^7^. Our data suggest immune hyperactivation can also induce lethality during development and that Pngl prevents immune hyperactivation by helping establish a normal gut barrier.

Although *foxo* heterozygosity had the biggest rescue effect in *Pngl* mutants, loss of one copy of *Rel* and *Tl* also showed some degree of lethality rescue. Since pathogens are thought to be the main inducers of the Imd and Toll pathways in *Drosophila*^64^, we anticipated that gut microbiota contribute to the lethality of *Pngl^–/–^* animals. Rather surprisingly, germ-free rearing of *Pngl* mutants did not rescue lethality at all, even though it dramatically rescued the developmental delay in these animals. How can one reconcile these observations? Since germ-free rearing also removes commensal bacteria, one possibility is that commensal bacteria are important for the gut homeostasis in *Pngl* mutants. Regardless of the reason for this discrepancy, our data indicate that upon gut barrier defect in *Pngl* mutants, infectious insults are not the primary mediators of hyperactive immune response.

Mammalian NGLY1 is shown to depend on its deglycosylation activity to mediate its biological functions^29,49^. Initially, it was thought that the *Drosophila* Pngl might lack deglycosylation activity^24^. However, it was later demonstrated that the fly Pngl has a deglycosylation activity comparable to human NGLY1 and that a point mutation in the catalytic domain of the fly Pngl fully abolished its ability to rescue the lethality of *Pngl^–/–^* animals^25^. About one third of the proteome in animals is thought to be decorated with *N-* glycans^65^. Therefore, theoretically, thousands of proteins can be potential targets for NGLY1 and its homologs. To date, two animal proteins have been shown to depend on the deglycosylation activity of NGLY1 for their function, SKN-1/NFE2L1 and Dpp/BMP4^29,31,49^. We have previously shown that Pngl regulates Dpp in the visceral mesoderm, not endoderm^25^. Moreover, our current data suggest that impaired NFE2L1 activity cannot explain the hyperactivation of innate immune response genes in *Pngl* mutant midguts (Supplementary Fig. 1). Therefore, the direct targets of Pngl mediating the phenotypes studied in this work remain to be identified. Accumulation of ConA^+^ and WGA^+^ puncta in PM-secreting cells in *Pngl^–/–^* larvae suggests that loss of Pngl affects the secretion of some *N*-glycoproteins. Components of the *Drosophila* larval PM have not been systematically identified. However, studies in other insects indicate that some PM components are *N*-glycoproteins^16,17^. Therefore, the gut barrier defects observed in *Pngl^–/–^* larvae might result from a failure in the deglycosylation of key *N*-glycoprotein components of the PM. Human insulin receptors harbors 19 predicted *N-*glycosylation sites, 14 of which have been experimentally verified^66^. Importantly, *Drosophila* InR is predicted to have 12 *N*-glycosylation sites (*NetNGlyc 1.0 server prediction*). Therefore, while the reduction in InR signaling in *Pngl^–/–^* larvae can be explained by starvation and reduced expression of *InR*, it is also plausible that the InR protein itself is a direct target of Pngl and shows abnormal trafficking and/or function upon loss of Pngl-mediated deglycosylation. Further studies are required to identify and characterize the biologically relevant targets of NGLY1/Pngl in the *Drosophila* larval intestine and fat body.

Human patients with NGLY1 deficiency display an array of symptoms including global developmental delay, lack of tears, and chronic constipation^67^. Some of the NGLY1 deficiency patients were reported to have recurrent, severe respiratory infections, while others were reported to have higher than expected antibody titers against rubella and rubeola after Measles, Mumps and Rubella (MMR) vaccination^22^. Moreover, global gene profiling in *Ngly1*-deficient melanoma cells showed upregulation of cytokines such as interferon β1 and interleukin 29 (ref. ^68^), and *Ngly1^–/–^* mouse embryonic fibroblasts exhibit increased expression of the interferon genes^30^. These observations suggest an association between loss of NGLY1 and altered immune response in various contexts. Moreover, altered glycosylation in intestinal epithelial cells is implicated in chronic inflammatory diseases including inflammatory bowel disease^69^. The critical roles uncovered here for *Drosophila* Pngl in regulating the gut mucus barrier, innate immune response, and metabolic homeostasis warrant further studies to explore whether loss of NGLY1 in other systems or alterations in *N*-glycosylation machinery exert similar immune and metabolic effects.

## Methods

### *Drosophila* strains and culture

Animals were reared at room temperature on standard food containing cornmeal, molasses, and yeast in all experiments except for those involving high-protein, intermediate-fat diet (HPIFD) and high-fat diet (HFD). Detailed composition of these diets is listed in Supplementary Table 2. The following *Drosophila* strains were used in the study: (1) *y w*, (2) *foxo^Δ94^/TM6B, Tb^1^*, (3) *Tl/TM3, Sb^1^*, (4) *Rel^E38^*, (5) *UAS-foxo^RNAi^*, (6) *UAS-Tl^RNAi^*, (7) *y^1^ sc* v^1^ sev^21^;UAS-Rel^RNAi^*, (8) *Myo1A-GAL4*, (9) *r4-GAL4*, (10) *c135-GAL4*, (11) *UAS-dipl2*, (12) *UAS-InR^WT^*, (13) *UAS-InR^A1325D^* (constitutively activated *UAS-InR^CA^*), (14) *UAS-Akt^WT^*, (15) *UAS-Akt^ΔPH^* (constitutively activated *UAS-Akt^CA^*), and (16) *UAS-bsk^RNAi^* (Bloomington Drosophila Stock Center); (17) *Mef2-GAL4* (ref. ^70^); (18) *Pngl^ex14^* and (19) *UAS-Pngl^WT^* (ref. ^24^); (20) *PBac{Pngl^wt^}VK31* (*Pngl* duplication; ref. ^26^); and (21) *UAS-dipl6* (ref. ^71^).

### Transcriptomic analysis/RNA sequencing

Midgut tissue from third instar larvae was dissected and homogenized in group of 25 in cold solution of Tri-reagent (Sigma-Aldrich, T9424); the amount of RNA in each sample was determined by Nanodrop, and RNA quality was analyzed using agarose gel electrophoresis (1.2%). The samples were prepared on a Beckman FXP using the Illumina TruSeq stranded mRNA chemistry and sequenced on a NextSeq 500 (Mid Output flowcell) in paired-end mode. Raw paired-end sequencing reads were trimmed using cutadapt v1.12 to remove Illumina adapters and low-quality bases. The processed reads were then aligned to the Drosophila dm6 genome using STAR v2.5.3a. Gene-level counts, based on GTF annotations from Flybase Release 6.20, were tabulated by totaling all reads overlapping the collapsed set of exons for each gene following previously published methods^72^. Genes with a counts per million reads mapped (CPM) of less than 5 in less than two samples were excluded. Differentially expressed genes were identified by using the edgeR glmFit model. The false discovery rate (FDR) was controlled by applying the Benjamini–Hochberg procedure to the p-values. Significantly expressed genes were defined as those exhibiting an absolute fold-change of at least 1.5 and an FDR of less than or equal to 0.01. The raw data have been deposited in the Gene Expression Omnibus (GEO) under accession number GSE206229.

### Lethality rescue assay

Lethality rescue was examined according to previously published report^25^. We scored the number of eclosed progeny (observed). The estimated total number (expected) was calculated based on Mendelian inheritance for each genotype. The observed/expected ratio is presented as lethality rescue percentage.

### Real time quantitative RT-PCR analysis

Total RNA was extracted from 5 larval midguts with Trizol (Invitrogen) and dissolved in 25 μL of RNase-free water. cDNA was then synthesized from 1 μg total RNA using amfiRivert II cDNA Synthesis Master Mix (R5500, GenDEPOT), and qPCR was carried out using amfiSure qGreen Q-PCR Master Mix, Low ROX (Q5601, GenDEPOT). Expression levels were normalized to *Actin* (endogenous control). Relative gene expression was calculated as fold change using the 2^-ΔΔCt^ method. The oligonucleotides used to assess target genes expression are listed in Supplementary Table 3.

### Dissections, staining, Image acquisition and processing

Larval midgut and fat body tissues were dissected and fixed in 4% paraformaldehyde. Antibodies were rabbit anti-dFoxo 1:250 (ab195977, abcam) and rabbit anti-SAPK/JNK 1:500 (Cat No. 559309, Sigma-Aldrich). Lectin staining in the proventriculus region was performed using helix pomatia agglutinin (HPA), Alexa Fluor^TM^ 488 conjugate 1:1000 (cat No. L11271, Invitrogen), wheat germ agglutinin (WGA) CF®488A 1:1000 (Cat No. 29022, Biotium), and concanavalin A (ConA) CF®488A 1:1000 (Cat No. 29016, Biotium). Confocal images were acquired using a Leica TCS-SP8 microscope and processed with Amira5.2.2. Image quantifications were performed using ImageJ. All images were processed in Adobe Photoshop CC. Figures were assembled in Adobe Illustrator CC.

### Western blotting

Protein samples were prepared from larval midguts in lysis buffer containing Halt™ Phosphatase Inhibitor Single-Use Cocktail (Thermo Fisher Cat. No. 78428) and Protease Inhibitor Cocktail (Promega Cat. No. G6521). The following antibodies were used: rabbit anti-pFoxo1 1:1000 (Cat No. 9461, Cell Signaling Technology), rabbit anti-Foxo1 1:1000 (Cat No. 2880, Cell Signaling Technology), rabbit anti-Akt 1:1000 (Cat No. 4691, Cell Signaling Technology), rabbit anti-pAkt 1:1000 (cat No. 4060, Cell Signaling technology) and mouse anti-actin 1:1000 (DSHB Cat. No. 224236-1). Western blots were developed using Clarify ECL Western Blotting Substrates (BioRad). The bands were detected using an Azure Biosystems c280 digital imager using chemiluminescent detection of HRP. Three independent immunoblots were performed for each experiment.

### Feeding behavior assay

Larval feeding behavior method was adapted from ref. ^73^. Third instar larvae were placed in sucrose-agar plates (5% sucrose mixed in 3% agar medium) and allowed to settle for 15 min. Number of mouth-hook contractions per minute were counted. A total number of 30 larvae were scored for each group.

### Generation of germ-free animals

Germ-free animals were generated by following the previously published method^74^. Flies were kept for egg laying in grape juice agar plate for 3-4 hr. Eggs were collected and dechorionated with 2.7% sodium hypochlorite for 2-3 minutes. Dechorionated eggs were washed twice in 70% ethanol and thrice in water, and then transferred to sterile fly food containing tetracycline (50 µg/mL). Flies were reared on sterile food vials for three generations. Absence of bacteria in germ-free flies was confirmed by 16S rRNA amplification using the 27F (5’-AGAGTTTGATCCTGGCTCAG-3’) and 1492R (5’ GGTTACCTTGTTACGACTT 3’) primers.

### Developmental assay

For developmental assay, the expected ratio was calculated based on Mendelian inheritance for genotypic classes and the observed/expected ratio is reported as a percentage. Crosses made up of 5 virgin females and 5 males were set in each tube after a period for sexual maturation and housing to obtain the maximum fitness level during the three days of egg deposition in a non-overcrowded environment. Crosses were set in triplicate and flies were transferred to fresh vials every 3 days for three times. The total number of pupae for genotypic classes produced over 20 days in each vial was scored.

### Nile red staining

Midgut and fat body tissues from third instar larvae were dissected and fixed in 4% paraformaldehyde. Fixation and washing of tissues were followed by the incubation in with 1:2500 dilution of 0.5mg/mL Nile red (Cat No. 19123, Sigma Aldrich) for half an hour. Upon incubation, tissues were rinsed in water and mounted in 80% glycerol followed by image acquisition.

### Triacyglycerol and free fatty acid estimation

Triacylglycerol and free fatty acid levels were estimated from larval midgut and hemolymph. Midgut samples were prepared by homogenizing ∼15 midguts in cold 1X PBST (0.05% tween). Hemolymph was collected from 15 larvae and mixed in cold 1X PBST. Triacylglycerol levels were estimated using manufacturer’s protocol (Sigma Aldrich # MAK266). Free fatty acid levels were estimated using manufacturer’s protocol (Sigma Aldrich # MAK044). Triacylglycerol and free fatty acid levels were normalized with the protein content determined by Bradford assay.

### Dextran feeding assay and gut barrier defect quantification

Third instar larvae were fed on semi-solid drops of 500 kDa FITC-labelled dextran (Sigma-Aldrich, St. Louis, MO, USA), diluted to 1 mg/mL in sucrose-agar medium (5% sucrose mixed in 3% agar medium) on a petri dish. Larvae were allowed to feed for 15 minutes. Larvae were washed with cold 1X PBS to remove any excess FITC-dextran on the surface. Midguts were dissected out and fixed in 4% paraformaldehyde followed by imaging under fluorescent microscope. For gut barrier defect quantification, larval guts that failed to retain FITC-signal (green) in their lumen were scored. Pharmacological induction of gut barrier defects was achieved by rearing third instar larvae on food containing 100 µM polyoxin D (Sigma Aldrich # 529313) for 48 hr. Gut barrier defects were examined and quantified using the dextran feeding assay described above.

### Statistical analysis

Unpaired Student’s t test, one-way ANOVA and two-way ANOVA with the Dunett’ and Bonferroni Post hoc tests, respectively, were used for statistical analyses. Statistical tests, *P* values and parameters including the sample sizes and replicates are mentioned in Figure Legends. All statistical analyses were performed using GraphPad Prism 9.

### Data Availability

Data are available with in the article and supplementary information. RNAseq data set are available through the Gene Expression Omnibus (GEO) accession number GSE206229.

## Acknowledgements

We acknowledge support from the NIH (R35GM130317 to H.J.N.), the Mizutani Foundation for Glycoscience (grant #220061 to H.J.N.), the Grace Science Foundation (research grants to H.J.N. and L.M.S.), AIRC (Investigator grant 20661 to T.V) and MUR (PRIN investigator grant 2020CLZ5XW to T.V.). We thank the Bloomington *Drosophila* Stock Center (NIH P40OD018537) and the Developmental Studies Hybridoma Bank for reagents; the Microscopy Core of the Baylor College of Medicine (BCM) Intellectual and Developmental Disabilities Research Center (IDDRC) (supported by 1U54HD083092 from the Eunice Kennedy Shriver National Institute of Child Health and Human Development [NICHD/NIH]), the BCM Integrated Microscopy Core (supported by NCI-CA125123, NIDDK-56338-13/ 15, and CPRIT-RP150578 and the John S. Dunn Gulf Coast Consortium for Chemical Genomics), Debra Townley for assistance with sectioning midguts for light microscopy, and Professor Pierre Leopold for generously providing *UAS-dilp6* strain and *Drosophila* anti-Foxo antibody. We also thank members of the Jafar-Nejad lab for discussions.

## Authors contribution

A.P., A.G., S.Y.H, T.V. and H.J.-N. designed and conceived the project and interpreted the data. W.F.M., B.S. and L.M.S. performed RNAseq and transcriptomic analysis. A.P., A.G., S.Y.H., G.C. performed the experiments. A.P., A.G., T.V. and H.J.-N. wrote the manuscript. All authors read, edited and approved the manuscript.

## References

1. Kvidera, S.K. et al. Intentionally induced intestinal barrier dysfunction causes inflammation, affects metabolism, and reduces productivity in lactating Holstein cows. J Dairy Sci 100, 4113–4127 (2017).

2. Pan, Z., Yuan, X., Tu, W., Fu, Z. & Jin, Y. Subchronic exposure of environmentally relevant concentrations of F-53B in mice resulted in gut barrier dysfunction and colonic inflammation in a sex-independent manner. Environ Pollut 253, 268–277 (2019).

3. Rera, M., Clark, R.I. & Walker, D.W. Intestinal barrier dysfunction links metabolic and inflammatory markers of aging to death in Drosophila. Proc Natl Acad Sci U S A 109, 21528–21533 (2012).

4. Zhu, R. & Ma, X.C. Role of metabolic changes of mucosal layer in the intestinal barrier dysfunction following trauma/hemorrhagic shock. Pathol Res Pract 214, 1879–1884 (2018).

5. Takiishi, T., Fenero, C.I.M. & Camara, N.O.S. Intestinal barrier and gut microbiota: Shaping our immune responses throughout life. Tissue Barriers 5, e1373208 (2017).

6. Blach-Olszewska, Z. & Leszek, J. Mechanisms of over-activated innate immune system regulation in autoimmune and neurodegenerative disorders. Neuropsychiatr Dis Treat 3, 365–372 (2007).

7. Badinloo, M. et al. Overexpression of antimicrobial peptides contributes to aging through cytotoxic effects in Drosophila tissues. Arch Insect Biochem Physiol 98, e21464 (2018).

8. Cao, Y., Chtarbanova, S., Petersen, A.J. & Ganetzky, B. Dnr1 mutations cause neurodegeneration in Drosophila by activating the innate immune response in the brain. Proc Natl Acad Sci U S A 110, E1752–1760 (2013).

9. Kominsky, D.J., Campbell, E.L. & Colgan, S.P. Metabolic shifts in immunity and inflammation. J Immunol 184, 4062–4068 (2010).

10. Hotamisligil, G.S. Inflammation, metaflammation and immunometabolic disorders. Nature 542, 177–185 (2017).

11. Zhao, L. et al. Chronic inflammation aggravates metabolic disorders of hepatic fatty acids in high-fat diet-induced obese mice. Sci Rep 5, 10222 (2015).

12. Paone, P. & Cani, P.D. Mucus barrier, mucins and gut microbiota: the expected slimy partners? Gut 69, 2232–2243 (2020).

13. Lehane, M.J. Peritrophic matrix structure and function. Annu Rev Entomol 42, 525–550 (1997).

14. Rizki, M.T.M. The secretory activity of the proventriculus of Drosophila melanogaster. J Exp Zool 131, 203–221 (1956).

15. Lehane, M.J., Allingham, P.G. & Weglicki, P. Composition of the peritrophic matrix of the tsetse fly, Glossina morsitans morsitans. Cell Tissue Res 283, 375–384 (1996).

16. Liu, D. et al. The N-glycan profile of the peritrophic membrane in the Colorado potato beetle larva (Leptinotarsa decemlineata). J Insect Physiol 115, 27–32 (2019).

17. Eisemann, C.H. et al. Larvicidal activity of lectins on Lucilia cuprina: mechanism of action. Entomol Exp Appl 72, 1–10 (1994).

18. Suzuki, T., Park, H. & Lennarz, W.J. Cytoplasmic peptide:N-glycanase (PNGase) in eukaryotic cells: occurrence, primary structure, and potential functions. FASEB J 16, 635–641 (2002).

19. Hosomi, A., Fujita, M., Tomioka, A., Kaji, H. & Suzuki, T. Identification of PNGase-dependent ERAD substrates in Saccharomyces cerevisiae. Biochem J 473, 3001-3012 (2016).

20. Hirsch, C., Blom, D. & Ploegh, H.L. A role for N-glycanase in the cytosolic turnover of glycoproteins. EMBO J 22, 1036–1046 (2003).

21. Enns, G.M. et al. Mutations in NGLY1 cause an inherited disorder of the endoplasmic reticulum-associated degradation pathway. Genet Med 16, 751–758 (2014).

22. Lam, C. et al. Prospective phenotyping of NGLY1-CDDG, the first congenital disorder of deglycosylation. Genet Med 19, 160–168 (2017).

23. Need, A.C. et al. Clinical application of exome sequencing in undiagnosed genetic conditions. J Med Genet 49, 353–361 (2012).

24. Funakoshi, Y. et al. Evidence for an essential deglycosylation-independent activity of PNGase in Drosophila melanogaster. PLoS One 5, e10545 (2010).

25. Galeone, A. et al. Tissue-specific regulation of BMP signaling by Drosophila N-glycanase 1. Elife 6, e27612 (2017).

26. Han, S.Y. et al. A conserved role for AMP-activated protein kinase in NGLY1 deficiency. PLoS Genet 16, e1009258 (2020).

27. Huang, D.W., Sherman, B.T. & Lempicki, R.A. Systematic and integrative analysis of large gene lists using DAVID bioinformatics resources. Nat Protoc 4, 44–57 (2009).

28. Sherman, B.T., et al. DAVID: a web server for functional enrichment analysis and functional annotation of gene lists (2021 update). Nucleic Acids Res 50, W216–W221 (2022).

29. Tomlin, F.M. et al. Inhibition of NGLY1 Inactivates the Transcription Factor Nrf1 and Potentiates Proteasome Inhibitor Cytotoxicity. ACS Cent Sci 3, 1143–1155 (2017).

30. Yang, K., Huang, R., Fujihira, H., Suzuki, T. & Yan, N. N-glycanase NGLY1 regulates mitochondrial homeostasis and inflammation through NRF1. J Exp Med 215, 2600–2616 (2018).

31. Lehrbach, N.J. & Ruvkun, G. Proteasome dysfunction triggers activation of SKN-1A/Nrf1 by the aspartic protease DDI-1. eLife 5, e17721 (2016).

32. Owings, K.G., Lowry, J.B., Bi, Y., Might, M. & Chow, C.Y. Transcriptome and functional analysis in a Drosophila model of NGLY1 deficiency provides insight into therapeutic approaches. Human molecular genetics 27, 1055–1066 (2018).

33. Rauscher, B. et al. Patient-derived gene and protein expression signatures of NGLY1 deficiency. J Biochem 171, 187–199 (2022).

34. Tzou, P. et al. Tissue-specific inducible expression of antimicrobial peptide genes in Drosophila surface epithelia. Immunity 13, 737–748 (2000).

35. Bulet, P., Hetru, C., Dimarcq, J.L. & Hoffmann, D. Antimicrobial peptides in insects; structure and function. Dev Comp Immunol 23, 329–344 (1999).

36. Dinkova-Kostova, A.T., Fahey, J.W., Kostov, R.V. & Kensler, T.W. KEAP1 and Done? Targeting the NRF2 Pathway with Sulforaphane. Trends in food science & technology 69, 257–269 (2017).

37. Buchon, N., Silverman, N. & Cherry, S. Immunity in Drosophila melanogaster--from microbial recognition to whole-organism physiology. Nat Rev Immunol 14, 796–810 (2014).

38. Guo, L., Karpac, J., Tran, S.L. & Jasper, H. PGRP-SC2 promotes gut immune homeostasis to limit commensal dysbiosis and extend lifespan. Cell 156, 109–122 (2014).

39. Becker, T. et al. FOXO-dependent regulation of innate immune homeostasis. Nature 463, 369–373 (2010).

40. Fink, C. et al. Intestinal FoxO signaling is required to survive oral infection in Drosophila. Mucosal Immunol 9, 927–936 (2016).

41. Hanson, M.A. et al. Synergy and remarkable specificity of antimicrobial peptides in vivo using a systematic knockout approach. Elife 8, e44341 (2019).

42. Eijkelenboom, A. & Burgering, B.M. FOXOs: signalling integrators for homeostasis maintenance. Nat Rev Mol Cell Biol 14, 83–97 (2013).

43. Puig, O., Marr, M.T., Ruhf, M.L. & Tjian, R. Control of cell number by Drosophila FOXO: downstream and feedback regulation of the insulin receptor pathway. Genes Dev 17, 2006–2020 (2003).

44. Junger, M.A. et al. The Drosophila forkhead transcription factor FOXO mediates the reduction in cell number associated with reduced insulin signaling. J Biol 2, 20 (2003).

45. Rodriguez, T.P. et al. Defects in the Neuroendocrine Axis Contribute to Global Development Delay in a Drosophila Model of NGLY1 Deficiency. G3 (Bethesda) 8, 2193–2204 (2018).

46. Rulifson, E.J., Kim, S.K. & Nusse, R. Ablation of insulin-producing neurons in flies: growth and diabetic phenotypes. Science 296, 1118–1120 (2002).

47. Okamoto, N. et al. A fat body-derived IGF-like peptide regulates postfeeding growth in Drosophila. Dev Cell 17, 885–891 (2009).

48. Blom, N., Sicheritz-Ponten, T., Gupta, R., Gammeltoft, S. & Brunak, S. Prediction of post-translational glycosylation and phosphorylation of proteins from the amino acid sequence. Proteomics 4, 1633–1649 (2004).

49. Galeone, A. et al. Regulation of BMP4/Dpp retrotranslocation and signaling by deglycosylation. Elife 9, e55596 (2020).

50. Britton, J.S., Lockwood, W.K., Li, L., Cohen, S.M. & Edgar, B.A. Drosophila’s insulin/PI3-kinase pathway coordinates cellular metabolism with nutritional conditions. Dev Cell 2, 239–249 (2002).

51. Zhang, L., Turner, B., Ribbeck, K. & Ten Hagen, K.G. Loss of the mucosal barrier alters the progenitor cell niche via Janus kinase/signal transducer and activator of transcription (JAK/STAT) signaling. J Biol Chem 292, 21231–21242 (2017).

52. Zhang, L. et al. O-glycosylation regulates polarized secretion by modulating Tango1 stability. Proc Natl Acad Sci U S A 111, 7296–7301 (2014).

53. Asahina, M. et al. JF1/B6F1 Ngly1(-/-) mouse as an isogenic animal model of NGLY1 deficiency. Proc Jpn Acad Ser B Phys Biol Sci 97, 89–102 (2021).

54. Yoshida, Y. et al. Loss of peptide:N-glycanase causes proteasome dysfunction mediated by a sugar-recognizing ubiquitin ligase. Proc Natl Acad Sci U S A 118, e2102902118 (2021).

55. Fujihira, H. et al. Liver-specific deletion of Ngly1 causes abnormal nuclear morphology and lipid metabolism under food stress. Biochim Biophys Acta Mol Basis Dis 1866, 165588 (2020).

56. Rudin, W. & Hecker, H. Lectin-binding sites in the midgut of the mosquitoes Anopheles stephensi Liston and Aedes aegypti L. (Diptera: Culicidae). Parasitol Res 75, 268–279 (1989).

57. Clark, R.I. et al. Distinct Shifts in Microbiota Composition during Drosophila Aging Impair Intestinal Function and Drive Mortality. Cell Rep 12, 1656–1667 (2015).

58. Rodgers, F.H., Gendrin, M., Wyer, C.A.S. & Christophides, G.K. Microbiota-induced peritrophic matrix regulates midgut homeostasis and prevents systemic infection of malaria vector mosquitoes. PLoS Pathog 13, e1006391 (2017).

59. Shahabuddin, M., Kaidoh, T., Aikawa, M. & Kaslow, D.C. Plasmodium gallinaceum: mosquito peritrophic matrix and the parasite-vector compatibility. Exp Parasitol 81, 386–393 (1995).

60. Biteau, B., Karpac, J., Hwangbo, D. & Jasper, H. Regulation of Drosophila lifespan by JNK signaling. Exp Gerontol 46, 349–354 (2011).

61. Wang, M.C., Bohmann, D. & Jasper, H. JNK extends life span and limits growth by antagonizing cellular and organism-wide responses to insulin signaling. Cell 121, 115–125 (2005).

62. Butterworth, F.M., Bodenstein, D. & King, R.C. Adipose Tissue of Drosophila Melanogaster. I. An Experimental Study of Larval Fat Body. J Exp Zool 158, 141–153 (1965).

63. Heier, C. & Kuhnlein, R.P. Triacylglycerol Metabolism in Drosophila melanogaster. Genetics 210, 1163–1184 (2018).

64. De Gregorio, E., Spellman, P.T., Tzou, P., Rubin, G.M. & Lemaitre, B. The Toll and Imd pathways are the major regulators of the immune response in Drosophila. EMBO J 21, 2568–2579 (2002).

65. Stanley, P., Taniguchi, N. & Aebi, M. N-Glycans, in Essentials of Glycobiology. (eds. A. Varki, et al.) 99–111 (Cold Spring Harbor Laboratory Press, Copyright 2015-2017 by The Consortium of Glycobiology Editors, La Jolla, California. All rights reserved., Cold Spring Harbor (NY); 2015).

66. Sparrow, L.G. et al. N-linked glycans of the human insulin receptor and their distribution over the crystal structure. Proteins 71, 426–439 (2008).

67. Pandey, A., Adams, J.M., Han, S.Y. & Jafar-Nejad, H. NGLY1 Deficiency, a Congenital Disorder of Deglycosylation: From Disease Gene Function to Pathophysiology. Cells 11, 1155 (2022).

68. Zolekar, A. et al. Stress and interferon signalling-mediated apoptosis contributes to pleiotropic anticancer responses induced by targeting NGLY1. Br J Cancer 119, 1538–1551 (2018).

69. Reily, C., Stewart, T.J., Renfrow, M.B. & Novak, J. Glycosylation in health and disease. Nat Rev Nephrol 15, 346–366 (2019).

70. Brand, A.H. & Perrimon, N. Targeted gene expression as a means of altering cell fates and generating dominant phenotypes. Development 118, 401–415 (1993).

71. Slaidina, M., Delanoue, R., Gronke, S., Partridge, L. & Leopold, P. A Drosophila insulin-like peptide promotes growth during nonfeeding states. Dev Cell 17, 874–884 (2009).

72. Story, B. et al. Defining the expression of piRNA and transposable elements in Drosophila ovarian germline stem cells and somatic support cells. Life Sci Alliance 2, e201800211 (2019).

73. Shen, P. Analysis of feeding behavior of Drosophila larvae on solid food. Cold Spring Harb Protoc 2012, pdb.prot069328 (2012).

74. Ryu, J.H. et al. Innate immune homeostasis by the homeobox gene caudal and commensal-gut mutualism in Drosophila. Science 319, 777–782 (2008).

