## Supplementary Data for "Gut barrier defects, increased intestinal innate immune response, and enhanced lipid catabolism drive lethality in *N*-glycanase 1 deficient *Drosophila*"

Supplementary Figures 1-4

Supplementary Tables 1-3

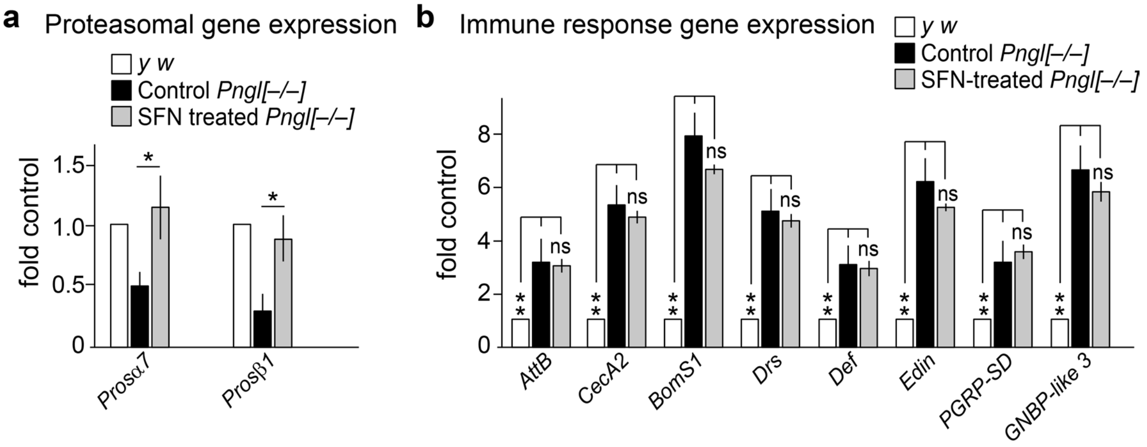


**Supplementary Figure 1. Related to Figure 1**

**Hyperactivation of immune genes in *Pngl* mutants cannot be explained by NFE2L1 associated proteasomal inhibition. a** Graph showing expression of proteasomal genes in the indicated genotypes. Error bars represent SD of three replicates. Significance is ascribed as *$P$<0.05 using one-way ANOVA. **b** Graph showing expression of immune genes in the indicated genotypes. Error bars represent SD of three replicates. Significance is ascribed as **$P$<0.01 using one-way ANOVA. ns, not significant.

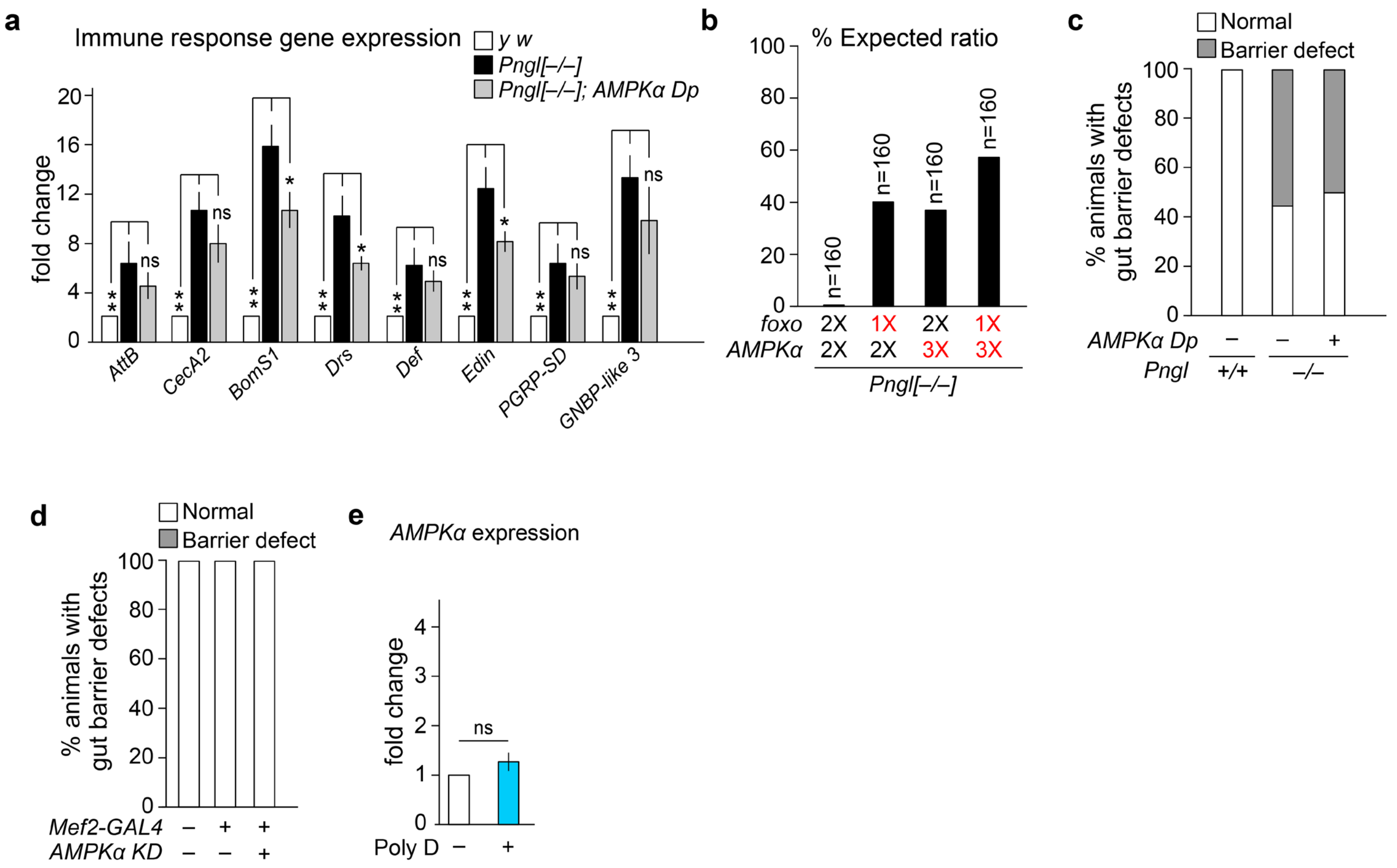


**Supplementary Figure 2. Related to Figure 1**

**Innate immune gene activation and gut barrier defect in *Pngl* mutants cannot be explained by decreased *AMPKα* level. a** Graph showing innate immune gene expression in the indicated genotypes. Error bars represent SD of three replicates. Significance is ascribed as **P*<0.05 and ***P*<0.01 using one-way ANOVA. ns, not significant. **b** Percentage lethality rescue in the indicated genotypes. n indicates the number of animals scored. **c** Graph showing quantification of the gut barrier defect phenotype in indicated genotypes (n=60 for each genotype). **d** Graph showing quantification of the gut barrier defect phenotype in indicated genotypes (n=60 for each genotype). **e** Graph showing *AMPKα* expression in Poly D-fed and control midguts of *y w* larvae. Error bars represent SD of three replicates. Significance is ascribed as **P*<0.05 and ***P*<0.01 using unpaired student's t-test. ns, not significant.

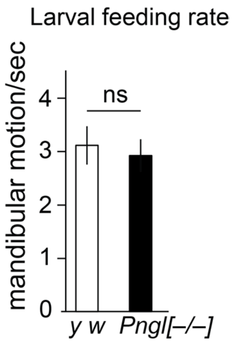


**Supplementary Figure 3. Related to Figure 6**

***Pngl* mutants exhibit feeding behavior comparable to wild-type.** Graph showing larval feeding behavior in wild-type and *Pngl* mutants (n=25 for each group). Error bars represent SD. Significance is ascribed as *$P$<0.05 and **$P$<0.01 using unpaired student's t-test. ns, not significant.

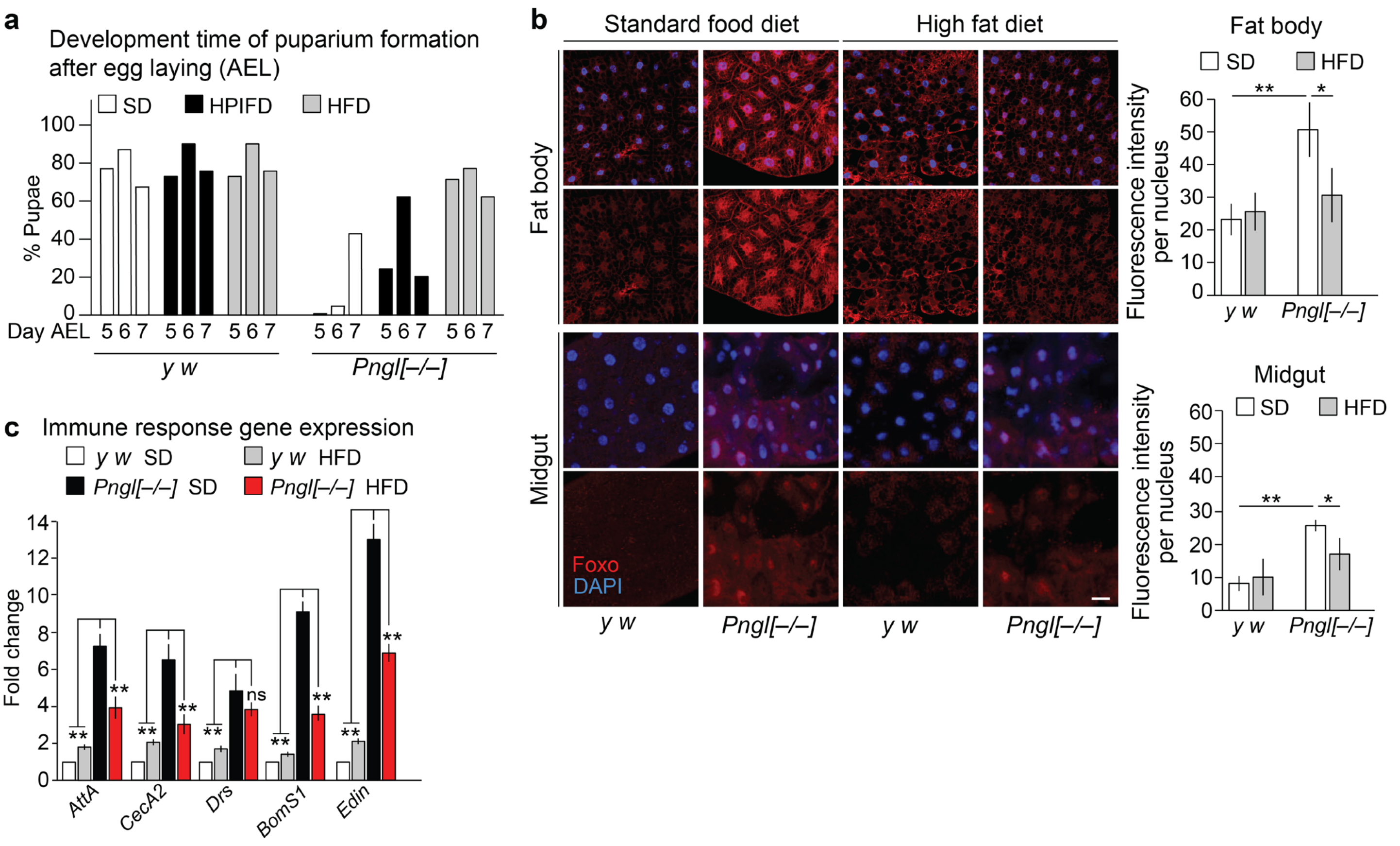


**Supplementary Figure 4. Related to Figure 7**

**HFD rescues the developmental delay and improves lethality rescue in *Pngl* mutants. a** Graph showing developmental time of puparium formation after egg laying (AEL) in control and *Pngl* mutants with the indicated diets. SD, standard diet; HPIFD, high-protein intermediate-fat diet; HFD, high-fat diet. **b** Confocal images showing DAPI (blue) and Foxo staining (red) and quantification of Foxo nuclear localization in the midgut and fat body of wild-type and *Pngl* mutants with the indicated diets. Scale bar is 50 µm. Error bars represent SD of three replicates. Significance is ascribed as *$P$<0.05 and **$P$<0.01 using two-way ANOVA. **c** Graph showing innate immune gene expression in control and *Pngl* mutants with or without HFD feeding. Error bars represent SD of three replicates. Significance is ascribed as *$P$<0.05 and **$P$<0.01 using two-way ANOVA. ns, not significant

**Supplementary Table 1. Related to Figure 1)**

**List of the genes differentially expressed in *Pngl*$^{-/-}$ midguts that exhibited >1.5 fold-change in all three pair-wise comparisons based on RNA-seq.** Provided as an Excel file.

**Supplementary Table 2. Related to Figure 7**

**Detailed composition of the standard diet (SD), high-protein, intermediate-fat diet (HPIFD), and high-fat diet (HFD) used in the study.**

| **Standard diet (SD)** | | | | | |
|---|---|---|---|---|---|
| Ingredients | Weight (g) | Calorie (kcal) | Fats (g) | Carbohydrates (g) | Proteins (g) |
| Water | 1000 | 0 | 0 | 0 | 0 |
| Yeast | 25 | 97.5 | 1.5 | 8.25 | 12.5 |
| Agar | 7 | 0 | 0 | 0 | 0 |
| Corn flour | 93 | 321.78 | 0.65 | 72.54 | 6.04 |
| Molasses | 155 | 444 | 0 | 118.4 | 0 |
| Propionic acid | 7 | 0 | 0 | 0 | 0 |
| Tegosept | 20 | 0 | 0 | 0 | 0 |

| **High-protein, intermediate-fat diet (HPIFD)** | | | | | |
|---|---|---|---|---|---|
| Ingredients | Weight (g) | Calorie (kcal) | Fats (g) | Carbohydrates (g) | Proteins (g) |
| Water | 1000 | 0 | 0 | 0 | 0 |
| Yeast | 19.23 | 75 | 1.15 | 6.35 | 9.62 |
| Agar | 7 | 0 | 0 | 0 | 0 |
| Corn flour | 19.23 | 67.31 | 0.14 | 15.15 | 1.26 |
| Sucrose | 32.3 | 129.23 | 0 | 32.3 | 0 |
| Propionic acid | 7 | 0 | 0 | 0 | 0 |
| Tegospot | 20 | 0 | 0 | 0 | 0 |
| Clarified butter | 25.92 | 232.77 | 25.85 | 0 | 0 |
| Soy protein | 69.6 | 258.31 | 1.04 | 0 | 62.65 |

| **High-fat diet (HFD)** | | | | | |
|---|---|---|---|---|---|
| Ingredients | Weight (g) | Calorie (kcal) | Fats (g) | Carbohydrates (g) | Proteins (g) |
| Water | 1000 | 0 | 0 | 0 | 00 |
| Yeast | 32.76 | 128.27 | 1.95 | 10.85 | 16.44 |
| Agar | 7 | 0 | 0 | 0 | 0 |
| Corn flour | 15.12 | 52.1 | 0.11 | 11.72 | 0.98 |
| Sucrose | 25.2 | 119.2 | 0 | 25.07 | 0 |
| Propionic acid | 7 | 0 | 0 | 0 | 0 |
| Tegospot | 20 | 0 | 0 | 0 | 0 |
| Clarified butter | 50.4 | 452.58 | 50.3 | 0 | 0 |

**Supplementary Table 3. List of the primer sequences used in the gene expression analysis in this study.**

| Gene | Primer (5'-3') |
|---|---|
| *AttA* | Forward- CTCCTGCTGGAAAACATC<br>Reverse- GCTCGTTTGGATCTGACC |
| *AttB* | Forward- GGGTAATATTTAACCGAAGT<br>Reverse-GTGCTAATCTCTGGTCATC |
| *CecA1* | Forward- CATTGGACAATCGGAAGCTGGGTG<br>Reverse- TAATCATCGTGGTCAACCTCGGGC |
| *CecA2* | Forward- ATTAGATAGTCATCGTGGTT<br>Reverse- GTGTTGGTCAGCACACT |
| *Def* | Forward- GTTCTTCGTTCTCGTGG<br>Reverse- CTTTGAACCCCTTGGC |
| *Drs* | Forward- AGTACTTGTTCGCCCTCTTGGCTG<br>Reverse- CCTTGTATCTTCCGGACAGGCAGT |
| *Edin* | Forward- AGTTCCAGACCAGTCCAGAG<br>Reverse- CGACCCACTTGGTTGTCCTT |
| *BomS1* | Forward- CTCGGTCTGCTGGCTGTGGC<br>Reverse- CCGTGGACATTGCACACCC |
| *Dso2* | Forward- CTGTCTGAAGATCTGCGGCT<br>Reverse- TTGAATCAACGTGTGTCCGC |
| *PGRP-SD* | Forward- ACTTGGATCGGTTTGCTCATC<br>Reverse- AGGGAGTTTCCATGCTGTCTAT |
| *PGRP-LA* | Forward- CTAAGGTGACCAGAAGCCCG<br>Reverse- CGGCCAGTTCCGGATTTCTT |
| *PGRP-SB1* | Forward- CCGCAATTTCAGCGATATTGG<br>Reverse- GGGAGTTGATCCCTGGAG |
| *GNBP-like 3* | Forward- GTCAAGGTCAACTCACCGAAG<br>Reverse- CGTGAAAAGCGAATAGGGAAATG |
| *Lip3* | Forward- AAAACGGGTGAATCTTCCAACC<br>Reverse- CCAGCATATAGGCCAGGGA |
| *CG6271* | Forward- ATGGATGTCGCTGAGGAGTG<br>Reverse- TGAAGAGGTAGAAGTTCACGGG |
| *CG6277* | Forward- TTGCCGAACAGTGGATGGAAG<br>Reverse- AAAGGTAGAACTTTACGGGAACG |
| *CG8093* | Forward- CACCATTGTTAGGGGACACGG<br>Reverse- GTGCATTGTGAGGATGTAACCA |
| *InR* | Forward- AAGCGTGGGAAAATTAAGATGGA<br>Reverse-GGCTGTCAACTGCTTCTACTG |
| *Actin5C* | Forward- TTGTCTGGGCAAGAGGATCAG<br>Reverse- ACCACTCGCACTTGCACTTTC |
